# Compositional control of FUS condensate ageing through aggregation-prone interaction networks

**DOI:** 10.64898/2026.07.31.741205

**Authors:** Eduardo Pedraza, Óscar Rebato, Alejandro Feito, Francisco Gámez, Pablo Llombart, Andrés R. Tejedor, Rosana Collepardo-Guevara, Jorge R. Espinosa

## Abstract

Stress granules are multicomponent biomolecular condensates whose aberrant ageing has been implicated in numerous neurodegenerative diseases. Although their composition is known to influence condensate properties, the molecular principles linking composition to structural maturation remain poorly understood. Here, we perform equilibrium and non-equilibrium residue-resolution molecular dynamics simulations to determine how RNA and heterotypic protein interactions regulate the pathological hardening of multicomponent FUS-containing condensates inspired by stress-granule composition. We show that diverse compositional changes—including RNA concentration, heterotypic protein partitioning, interfacial enrichment of G3BP1, and charged peptide recruitment—reshape condensate organization through distinct molecular mechanisms. Despite these different modes of action, all converge on a common physical principle: modulation of the local clustering and persistence of contacts between low-complexity aromatic-rich kinked segments (LARKS) governs the nucleation and accumulation of long-lived intermolecular cross-*β*-sheet structures. Intermediate RNA concentrations enhance condensate density and promote LARKS contacts, whereas high RNA levels, heterotypic interactions, and interfacial coating reduce their availability and delay ageing. Our results establish a unified molecular framework linking condensate composition, internal organization and ageing. This framework provides mechanistic insight into the regulation of multicomponent condensate material properties and suggests general design principles for modulating their pathological aggregation.

## I. INTRODUCTION

Membraneless organelles, known as biomolecular condensates, have emerged as fundamental organizers of cellular architecture, enabling cells to compartmentalize biomolecules without the need for lipid membranes^1–3^. These dynamic assemblies form through liquid-liquid phase separation, allowing proteins and nucleic acids to reversibly concentrate in space and time in response to physiological and environmental cues^4–10^. Among the best-characterized biomolecular condensates are ribonucleoprotein (RNP) granules, which are composed primarily of RNA-binding proteins (RBPs) and RNA molecules and regulate diverse aspects of RNA metabolism, gene expression and cellular adaptation to stress^11–15^. This family includes processing bodies (P-bodies), germ granules, neuronal granules and stress granules (SGs)^11,16–19^. SGs assemble transiently in response to a broad range of cellular insults, including oxidative stress, heat shock, nutrient deprivation and viral infection, largely as a consequence of translational arrest and polysome disassembly, which increase the cytoplasmic pool of untranslated mRNAs available to recruit RBPs through multivalent interactions^11,12^. Although the molecular mechanisms driving stress granule assembly have been extensively investigated, considerably less is known about how condensate composition governs their internal molecular organization and material properties.

SGs are assembled by a complex network of multivalent RNA-binding proteins, among which G3BP1 and its paralog G3BP2 function as central nucleators that coordinate the recruitment of numerous client proteins and RNAs^28,29^. Additional proteins, including FUS, TIA1, CAPRIN1, hnRNPA1 and TDP-43, further stabilize these condensates through extensive protein–protein and protein–RNA interaction networks mediated by intrinsically disordered regions (IDRs) enriched in low-complexity and prion-like domains^4,12,14, 30–35^. These weak multivalent interactions maintain the liquid-like character of SGs while allowing rapid molecular exchange with the surrounding cytoplasm. However, condensates are not static materials. Over time, they can undergo ageing, progressively reorganizing into increasingly stable intermolecular interaction networks that give rise to gel-like or solid-like material states^36–42^. This transition has attracted considerable attention because it represents a hallmark between functional condensates and pathological protein aggregation. In particular, low-complexity aromatic-rich kinked segments (LARKS) embedded within IDRs can nucleate inter-molecular cross-*β* structures that stabilize protein assemblies and hinder condensate dissolution under physiological conditions^4,33^. Dysregulation of these interactions through disease-associated mutations or post-translational modifications promotes irreversible aggregation and has been directly implicated in amyotrophic lateral sclerosis (ALS), frontotemporal dementia (FTD) and other related neurodegenerative disorders^43–47^.

Increasing evidence suggests that condensate composition is a major determinant of their material properties and propensity to undergo pathological ageing. RNA can either promote or suppress condensate ageing depending on its concentration and molecular context^38,48,49^. Likewise, heterotypic proteins may either enhance or compete with homotypic interactions in forming multiphase condensates^50–52^, whereas proteins such as TDP-43 or HSP70 can localize preferentially at condensate interfaces, altering molecular organization and cross-*β*-sheet accumulation without necessarily changing the overall composition^50,53^. More generally, the coexistence of multiple proteins and nucleic acids generates highly cooperative interaction networks whose emergent behaviour cannot be easily predicted from the properties of the individual components alone^36,44,54,55^. Despite these observations, it remains unclear whether the diverse compositional perturbations known to regulate condensate ageing operate through distinct molecular mechanisms or instead converge onto a common microscopic principle governing multicomponent condensate ageing.

Addressing this question experimentally remains challenging. Although fluorescence microscopy, spectroscopy and biochemical approaches have transformed our understanding of condensate self-assembly, they provide only limited access to the transient residue-level interaction networks that determine condensate organization and their evolution over time^11,52,56–59^. In particular, directly resolving how protein–protein and protein– RNA interactions, at molecular or submolecular level, are spatially organized within multicomponent condensates, how these interaction networks are remodelled during ageing, and how they ultimately regulate the nucleation of intermolecular cross-*β* structures remains extremely difficult^51,60,61^. These challenges are further compounded by the highly dynamic and heterogeneous composition of stress granules, whose molecular content depends strongly on cell type, stress conditions and exposure time^62^. Consequently, establishing general mechanistic principles linking condensate composition to time-dependent variations in the material properties remains a major challenge.

Computer simulations provide a unique opportunity to bridge this gap by directly connecting molecular interactions with condensate phase behaviour across multiple spatial and temporal scales^53,63–67^. Here, we investigate how distinct compositional perturbations regulate the accumulation of inter-protein *β*-sheet structures in multicomponent stress granules. To this end, we combine residue-resolution coarse-grained molecular dynamics simulations using the Mpipi-Recharged model^68^ with a nonequilibrium ageing algorithm^37,69^ that mimics disorder-to-order transitions associated with intermolecular cross-*β*-sheet formation. By systematically varying RNA concentration, heterotypic protein composition, interfacial organization and charged peptide recruitment, we show that seemingly distinct mechanisms of condensate reorganization converge onto a common microscopic principle: the regulation of the clustering and persistence of contacts between aggregation-prone LARKS domains. Our results establish a unified molecular framework linking condensate composition, internal biomolecular organization and structural hardening, providing mechanistic insight into how cells regulate stress granules stability and architecture, as well as suggesting general design principles for modulating pathological liquid-to-solid transitions driven by cross-*β*-sheet accumulation.

## II. RESULTS

### A. RNA reorganizes the FUS interaction network and regulates LARKS availability and condensate compaction

Several experimental and computational studies have shown that RNA is a key regulator of phase separation in ribonucleoprotein condensates^71–79^. At low concentrations, RNA promotes condensate formation by establishing favourable heterotypic protein–RNA interactions, whereas excess RNA destabilizes condensates through a reentrant phase transition mechanism^80^. Reproducing this characteristic behaviour therefore represents an important benchmark for computational models of biomolecular condensates. We first evaluated whether the Mpipi-Recharged residue-resolution implicit solvent model^68^ (technical details on the model are provided in Section S1 of the Supplementary Material, hereafter SM) describes the phase behaviour of FUS-polyAdenosine (polyA) RNA condensates. Using Direct Coexistence (DC) simulations^81^ (Fig. 1a), we determined the temperature–density phase diagrams of pure FUS condensates and condensates containing polyA single-stranded RNAs of 200 nucleotides at the polyA:FUS molar ratio of 1:5 (blue circles; see Section S3 of the SM). Consistent with *in vitro* experimental observations^75^, low RNA concentration stabilizes condensate formation, shifting the coexistence curve towards higher densities and increasing the critical temperature (*T*_*c*_) by approximately 20 K relative to pure FUS condensates (black circles; Fig. 1a).

**FIG. 1.**
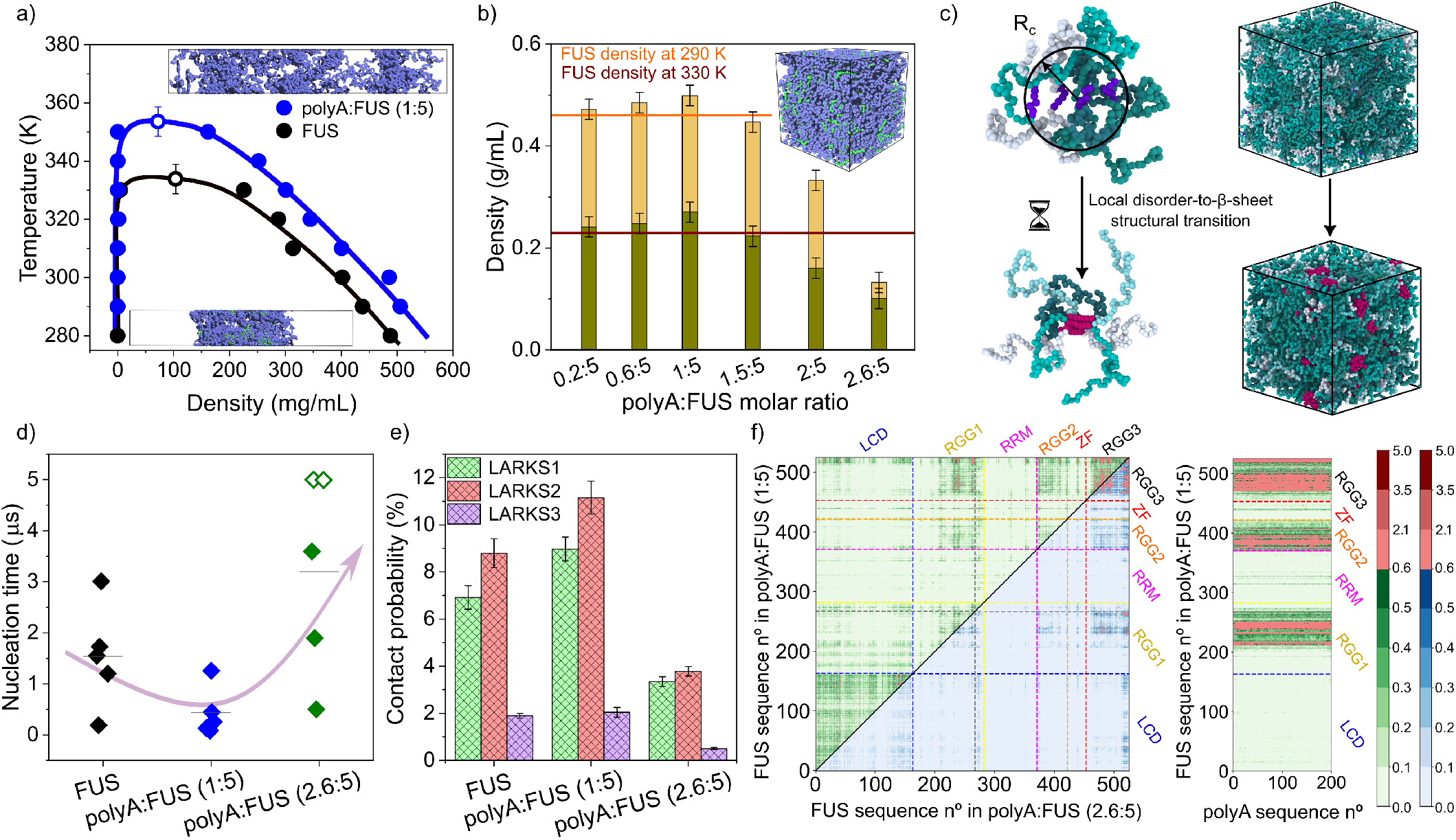
RNA induces non-monotonic condensate compaction and ageing in FUS-based condensates. (a) Temperature–density phase diagrams at NaCl concentration of 150 mM for FUS (black symbols) and a polyA:FUS mixture at molar ratio of 1:5 (blue symbols). Filled circles represent the coexistence densities obtained from DC simulations and empty symbols indicate the estimated critical points obtained through the law of rectilinear diameters and critical exponents^70^. (b) Density of polyA:FUS mixtures at different molar ratios evaluated at 290 and 330 K in orange and olive green, respectively. The horizontal lines account for the density of pure FUS condensates at the temperatures indicated in the legend. (c) Representation of the dynamic ageing algorithm used to model the disorder-to-order cross-*β*-sheet transitions. If four LARKS come within a distance smaller than the cut-off radius (*R*_*c*_), a structural transition is triggered, strengthening their protein-protein interactions. (d) Average nucleation time (⟨*τ*⟩, black horizontal bar) for the formation of FUS-associated *β*-sheets in pure FUS condensates and polyA:FUS mixtures with molar ratios of 1:5 and 2.6:5. Solid diamonds indicate nucleation times from individual trajectories where *β*-sheet transitions were observed. Empty diamonds correspond to condensates that did not undergo *β*-sheet formation within the 5 *µ*s simulation time (time at which are located to show a lower bound). (e) Contact probability of intermolecular LARKS–LARKS interactions in FUS (expressed as a percentage) at 290 K for condensates of FUS and polyA:FUS with molar ratios of 1:5 and 2.6:5. (f) Intermolecular contact maps (expressed in residue–residue percentage of contact frequency) for FUS in polyA:FUS condensates with molar ratios of 1:5 (upper diagonal) and 2.6:5 (lower diagonal) and for FUS–polyA in a polyA:FUS condensate with molar ratio of 1:5.

We next quantified the equilibrium density of polyA:FUS condensates over a larger range of RNA concentrations (Fig. 1b) using bulk *NpT* simulations as discussed in Ref.^82^. The model successfully reproduces the characteristic RNA-driven re-entrant phase behaviour observed experimentally for FUS^75^. Condensate density initially increases with RNA concentration, reaching a maximum with polyA:FUS molar ratio of 1:5 (see Fig. S2 for the equivalent mass-ratio representation), and subsequently decreases as additional RNA is incorporated into the solution. At this optimum composition, condensates reach equilibrium protein-RNA densities of approximately 0.50 g · mL^−1^ at 290 K and 0.27 g · mL^−1^ at 330 K, compared with 0.46 and 0.23 g · mL^−1^, respectively, for pure FUS condensates (water is excluded from condensate densities since the Mpipi-Recharged is an implicit solvent model). These results demonstrate that the Mpipi-Recharged reproduces the experimentally observed stabilization of FUS condensates at intermediate RNA concentrations, providing a robust foundation for investigating how RNA regulates the internal organization and ageing of FUS-scaffolded multicomponent stress granules.

We next investigated how polyA RNAs regulate the progressive accumulation of intermolecular cross*β* structures^37,41,83,84^ in FUS condensates. FUS contains across its low-complexity domain (LCD) three different LARKS^41^ (^37^SYSGYS^42, 54^SYSSYGQS^61^), and ^77^STGGYG^82^) susceptible to undergo disorder-to-order cross-*β*-sheet transitions. RNA has been proposed to either promote or suppress pathological condensate maturation depending on its concentration, length and molecular context^4,49,71,85^. To investigate the molecular origin of these observations, we combined simulations with a previously developed nonequilibrium local-dependent ageing algorithm^37,38,53,63^, which models the near irreversible disorder-to-order transition of FUS LARKS into inter-protein *β* structures (Fig. 1c). The algorithm introduces structural transitions only when neighbouring LARKS satisfy local geometric and cooperative criteria, allowing the progressive accumulation of fibrillar interactions while capturing the accompanying increase in protein–protein interactions and reduction in local conformational flexibility (see Refs.^37,38,86^ Section S5 of the SM for further details on the ageing algorithm).

We quantified the average nucleation time for the emergence of inter-protein *β*-sheet structures for five independent trajectories with distinct initial velocities (coloured diamonds) in bulk condensates at their equilibrium density containing different polyA:FUS molar ratios at 290 K (Fig. 1d). Remarkably, kinetic arrest driven by cross- *β*-sheet formation exhibits the same non-monotonic dependence on RNA concentration observed for condensate density and stability (Fig. 1b). Intermediate RNA concentrations markedly accelerate the nucleation of cross-*β* structures (blue diamonds), whereas higher RNA levels strongly delay their formation (green diamonds; where empty diamonds represent trajectories where no interprotein *β*-sheet transitions were observed within 5 *µ*s) compared to pure FUS condensates (black diamonds). This behaviour can be rationalized by the changes in condensate protein density induced by RNA. At intermediate concentrations, RNA promotes condensate compaction (Fig. 1b), increasing the local concentration of FUS LARKS motifs within its LCD, and therefore the probability of productive intermolecular structural transitions. In contrast, excess RNA induces condensate decompaction, reducing the local availability of LARKS clustering and substantially increasing the nucleation time. These results establish a direct connection between RNA-mediated condensate organization and the molecular kinetics of cross-*β*-sheet formation.

To uncover the molecular origin of the RNA-dependent ageing behaviour, we next analysed the intermolecular interaction networks within FUS and polyA:FUS condensates. In particular, we quantified the probability of intermolecular contacts involving the aggregation-prone LARKS motifs of FUS, which induce cross-*β* structures during condensate incubation^52,84^ (see Section S6 of the SM). Remarkably, the probability of LARKS–LARKS contacts closely follows the non-monotonic ageing kinetics observed in Fig. 1d. Relative to pure FUS condensates, LARKS contact probability increases substantially at a polyA:FUS molar ratio of 1:5 but decreases markedly at higher RNA concentrations (Fig. 1e). To understand the molecular basis of this behaviour, we examined the complete intermolecular contact map of FUS in contact percentage probability (Fig. 1f). At intermediate RNA concentrations, contacts between the FUS LCD and the RGG domains are strongly reduced relative to pure FUS condensates, whereas LCD–LCD interactions become significantly enriched. This reorganization arises because polyA binds preferentially to the positively charged RGG domains through electrostatic interactions, effectively competing with the cation–*π* interactions that normally stabilize LCD–RGG contacts (see Fig. S1 of the SM). As a consequence, the LCDs become increasingly available to establish homotypic interactions with neighbouring FUS molecules, thereby promoting encounters between LARKS motifs and facilitating the nucleation of cross-*β* structures. This mechanism is further reinforced by the increased compaction of condensates at intermediate RNA concentrations (Fig. 1b). Because a limited number of RNA molecules efficiently bridge multiple FUS proteins through favorable electrostatic interactions^78^, condensate density increases, further enhancing the local concentration of LARKS motifs and their probability of productive intermolecular encounters. In contrast, excess RNA reverses this behaviour. Additional RNA molecules saturate the RGG domains and increase electrostatic repulsion within the condensate, leading to condensate decompaction. Consequently, both the local concentration and persistence of FUS LCD–LCD interactions decrease, substantially reducing the availability of LARKS contacts required for cross-*β* nucleation (Fig. 1e). Together, these results identify RNA-mediated reorganization of intermolecular interaction networks as the molecular mechanism underlying the non-monotonic regulation of FUS condensates as a function of RNA concentration. Rather than acting directly on cross-*β* formation, RNA remodels the interaction network within the condensate, thereby regulating the local availability of LARKS contacts. This framework provides a molecular basis for understanding how compositional changes control condensate ageing and raises the question of whether this principle extends to other regulators of stress-granule organization.

### B. hnRNPA1 remodels FUS intermolecular interactions and delays condensate ageing

The molecular composition of biomolecular condensates determines the balance between homotypic and heterotypic interactions that govern their phase behaviour and internal organization^53,54,87^. In particular, previous studies have shown that the LCDs of FUS and hnRNPA1 readily form mixed condensates, where subtle changes in composition alter the thermodynamic driving forces underlying phase separation^88^. However, whether heterotypic protein mixing also regulates the nucleation of cross-*β*-sheet fibrils in FUS-hnRNPA1 condensates remains largely unknown, particularly in systems composed of the full-length proteins. To address this question, we performed DC simulations of condensates containing FUS mixed with: (1) full-length hnRNPA1, (2) the isolated hnRNPA1 LCD (A1+NLS), and (3) the phosphomimetic A1+12D variant^44^ (Fig. 2; see Sections S2 and S3 of the SM for further details on the sequences and simulations). We first quantified the degree of mixing between condensate components from their equilibrium density profiles (Fig. 2a,b and Fig. S3) and subsequently analysed how these compositional changes modify the intermolecular interaction network. In the previous section, we demonstrated that the probability of LARKS–LARKS contacts strongly correlates with the nucleation kinetics of cross-*β* structures (Figs. 1d,e), establishing this quantity as a microscopic descriptor of condensate ageing. We therefore use the local availability of LARKS–LARKS contacts to determine how heterotypic protein mixing modulates the propensity of condensates to undergo ageing.

**FIG. 2.**
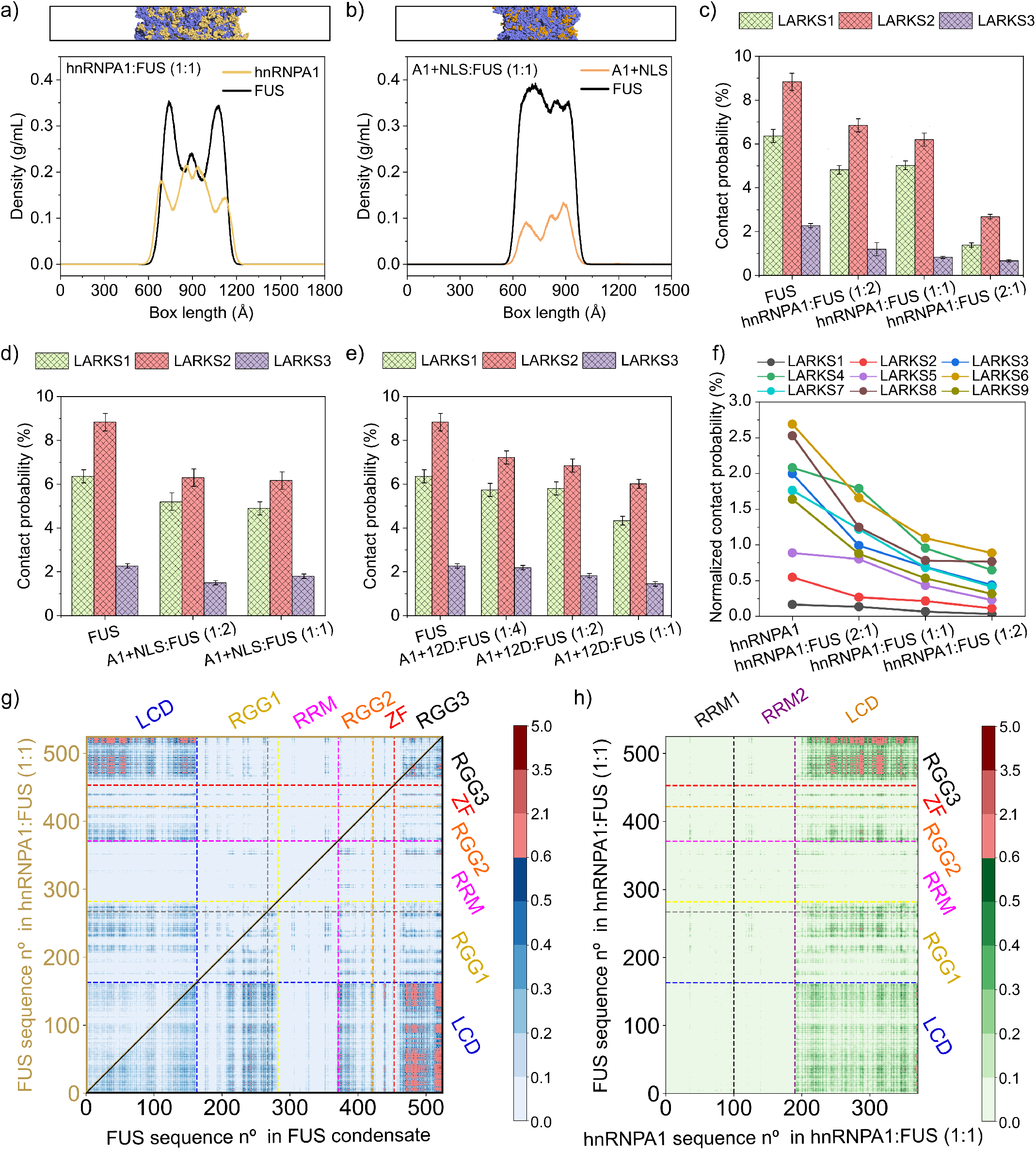
Protein scaffold mixing reduces homotypic LARKS contacts in heterotypic condensates. (a-b) Density profiles of condensates composed of hnRNPA1:FUS (a) and A1+NLS:FUS (b) mixtures at equimolar composition. Density profiles are shown for each component across the system. The *x* axis represents the length of the direct coexistence simulation box. (c-e) Contact probability of intermolecular LARKS–LARKS interactions among FUS replicas in condensates formed by hnRNPA1:FUS (c), A1+NLS:FUS (d), and A1+12D:FUS (e) mixtures with different molar ratios. (f) Normalized contact probability of intermolecular LARKS–LARKS interactions between hnRNPA1 in condensates composed by hnRNPA1:FUS mixtures with different molar compositions. The normalization accounts for the substantial differences. The normalization accounts for the substantial differences in the lengths of the identified hnRNPA1 segments (uncertainties are comparable to the size of the markers). (g) Intermolecular contact maps of FUS (expressed as residue–residue percentage of contact frequencies) in an equimolar hnRNPA1:FUS condensate (upper diagonal) and a pure FUS condensate (lower diagonal). (h) Intermolecular contact maps (expressed in % of residue contact frequency) of FUS–hnRNPA1 condensates at an equimolar concentration.

To determine whether the molecular principle identified for RNA also applies to heterotypic protein mixtures, we investigated how the incorporation of hnRNPA1 modifies the local availability of LARKS contacts within FUS condensates. We quantified these interactions from intermolecular contact maps obtained from DC simulations at 290 K (see Sections S3 and S6 of the SM). Across all condensate mixtures, increasing the fraction of hnRNPA1 consistently reduced the local availability of LARKS (Figs. 2c–e). In condensates formed by full-length hnRNPA1 and FUS, the probability of FUS LARKS–LARKS contacts decreased monotonically as the relative abundance of FUS was reduced (Fig. 2c). A similar trend was observed for mixtures containing the isolated hnRNPA1 LCD (A1+NLS) and the phosphomimetic A1+12D variant, although the reduction in LARKS contacts was less pronounced (see Figs. 2d,e and Fig. S3 of the SM). Interestingly, A1+12D:FUS condensates at low A1+12D fractions retained LARKS contact probabilities comparable to those of pure FUS condensates, indicating that the extent of LARKS suppression depends not only on condensate stoichiometry but also on the molecular sequence features of the interacting proteins. We also analyzed the equilibrium density profiles of the condensates (Figs. 2a,b and Fig. S3 of the SM). All systems exhibited a high degree of spatial mixing between FUS and hnRNPA1 species, demonstrating that heterotypic interactions dominate the condensate architecture. As the fraction of hnRNPA1 increases, heterotypic interactions increasingly compete with and dilute homotypic FUS–FUS contacts, weakening the network that brings FUS LARKS motifs into close proximity. This reduction in homotypic association lowers the local availability of aggregation-prone interactions and the probability of cross-*β* nucleation.

Having established that heterotypic protein mixing reduces the local availability of FUS LARKS, we next asked whether the same microscopic principle also applies to hnRNPA1, an RNA-binding protein also involved in stress granules formation^4,33,39^. Like FUS, hnRNPA1 undergoes structural ageing through the accumulation of intermolecular cross-*β* structures mediated by multiple LARKS motifs distributed throughout its sequence^60^. We therefore quantified the probability of intermolecular contacts between the reported hnRNPA1 LARKS (^187^ASASSSQ^193, 197^SGSGNF^202, 243^GYNGFG^248, 282^GGSGSYDSY^290, 301^GSGSNFG^307, 310^GSYNDF^315, 320^NQSSNF^325, 343^GGGQYF^348^, and ^358^GGSSSSSSYGS^368^) from DC simulations at increasing concentrations of FUS (Fig. 2f; see Section S2 of the SM). To account for differences in LARKS length, contact probabilities were normalized by the number of residues within each LARKS. Remarkably, hnRNPA1 exhibits the same qualitative behaviour as FUS. The local availability of hnRNPA1 LARKS contacts decreases monotonically as the molar fraction of hnRNPA1 in the condensate is reduced, indicating that heterotypic mixing suppresses homotypic aggregation-prone interactions in both proteins.

To uncover the molecular origin of the reduced local availability of LARKS-mediated contacts in mixed condensates, we next evaluated the intermolecular interaction networks underlying their structural organization. Fig. 2g compares the FUS–FUS intermolecular contact maps of a pure FUS condensate (below the diagonal) with those of a 1:1 hnRNPA1:FUS condensate (above the diagonal). Relative to the pure FUS system, the mixed condensate exhibits a pronounced reduction in both LCD– LCD and LCD–RGG interactions, demonstrating that recruitment of hnRNPA1 extensively remodels the homotypic interaction network responsible for FUS self-association. To identify the interactions replacing these contacts, we analyzed the heterotypic FUS–hnRNPA1 contact map and, specifically, which interactions increase as homotypic contacts diminish (Fig. 2h). The mixed condensate is dominated by extensive heterotypic interactions between the low-complexity domains of both proteins, together with a second interaction hotspot involving the hnRNPA1 LCD and the FUS RGG3 domain. These interaction patterns are fully consistent with the molecular grammar and composition encoded by each protein sequence (Section S2 and Fig. S1 of the SM). Both LCDs are highly enriched in aromatic residues, favouring extensive heterotypic *π*–*π* interactions, while the arginine residues distributed throughout the hnRNPA1 LCD establish additional cation–*π* interactions with aromatic residues within the FUS LCD. Rather than simply introducing new intermolecular contacts, hnRNPA1 therefore competes directly with FUS for the same interaction surfaces that normally stabilize homotypic FUS assemblies. This competitive rewiring of the intermolecular interaction network effectively diverts FUS molecules away from self-association, reducing both LCD–LCD contacts and the local availability of LARKS motifs required for cross-*β*-sheet formation.

Importantly, this mechanism is fundamentally different from that identified for polyA RNA. Whereas RNA remodels condensate organization by engaging the positively charged FUS RGG domains through favourable electrostatic protein–RNA interactions, hn-RNPA1 reorganizes the condensate through competitive heterotypic protein–protein interactions involving aromatic and cation–*π* contacts. Despite these distinct molecular pathways, both perturbations converge on the same microscopic outcome: a reduction in the local clustering of LARKS contacts at high stoichiometries. Together, these results support that structurally distinct interaction networks regulate condensate ageing through a common molecular principle, whereby condensate composition controls the probability of pathological maturation by modulating the frequency of productive encounters between aggregation-prone motifs.

### C. G3BP1 interfacial organization emerges as a regulator of FUS condensate ageing

So far, we have shown that condensate composition modulates ageing by controlling the degree of heterotypic mixing between its components and, consequently, the local availability of aggregation-prone contacts. However, the extent of molecular mixing represents only one aspect of condensate organization. Depending on the physicochemical properties of their constituents, multicomponent condensates can adopt markedly different spatial architectures, ranging from homogeneous mixtures to systems exhibiting pronounced interfacial enrichment or microphase segregation^52,87,89^. Such multiphasic architectures emerge from the interplay between interaction valency^54,81^, electrostatic complementarity^77,79,90^, and scaffold-mediated interactions^91,92^, which together determine how molecules are distributed within the condensed phase^36,88,93^. We therefore next asked whether the spatial organization of condensate components constitutes an additional mechanism regulating progressive kinetic arrest. To address this question, we investigated representative two-component condensates formed by FUS together with G3BP1 and FUS in presence of the positively charged peptide PR_25_, which may generate distinct condensate architectures through fundamentally different interaction networks.

To this end, we first characterized the internal architecture of G3BP1:FUS condensates at a (2.8:5) molar ratio using DC simulations. The average density profile of several trajectories spanning over 5 *mu*s (Fig. 3a), together with an equilibrium condensate simulation snapshot (top panel) and a primitive path analysis^38^ (PPA; bottom panel), revealed a strikingly heterogeneous organization in which G3BP1 forms an interfacial layer surrounding a FUS-rich core (see Section S7 of the SM). The LCD of FUS remains uniformly distributed throughout the FUS-rich phase (dashed black curve), indicating that G3BP1 selectively reorganizes the condensate at the mesoscale without inducing spatial segregation of the aggregation-prone regions within the FUS phase. We compared these results with condensates formed by FUS and the negatively charged A1+12D phosphomimetic variant (Fig. 3b) as well as with PR_25_ (Fig. 3c), an arginine-rich di-repeat peptide associated to ALS^94–96^. In both A1+12D:FUS and PR_25_:FUS equimolar condensate mixtures (Figs. 3b and c), the two components exhibited nearly homogeneous density distributions throughout the condensed phase, and the spatial distribution of the FUS LCD closely followed that of the full-length protein. Thus, unlike G3BP1, these sequences do not induce multiphasic condensate architectures. These results evidence that condensate composition determines not only the extent of heterotypic mixing but also the emergence of potential distinct spatial architectures, ranging from homogeneous mixtures to interfacially organized condensates depending on the specific sequence features^52^. These architectures can provide different structural environments in which aggregation-prone interaction networks emerge, and therefore may represent an additional level of regulation of condensate ageing.

**FIG. 3.**
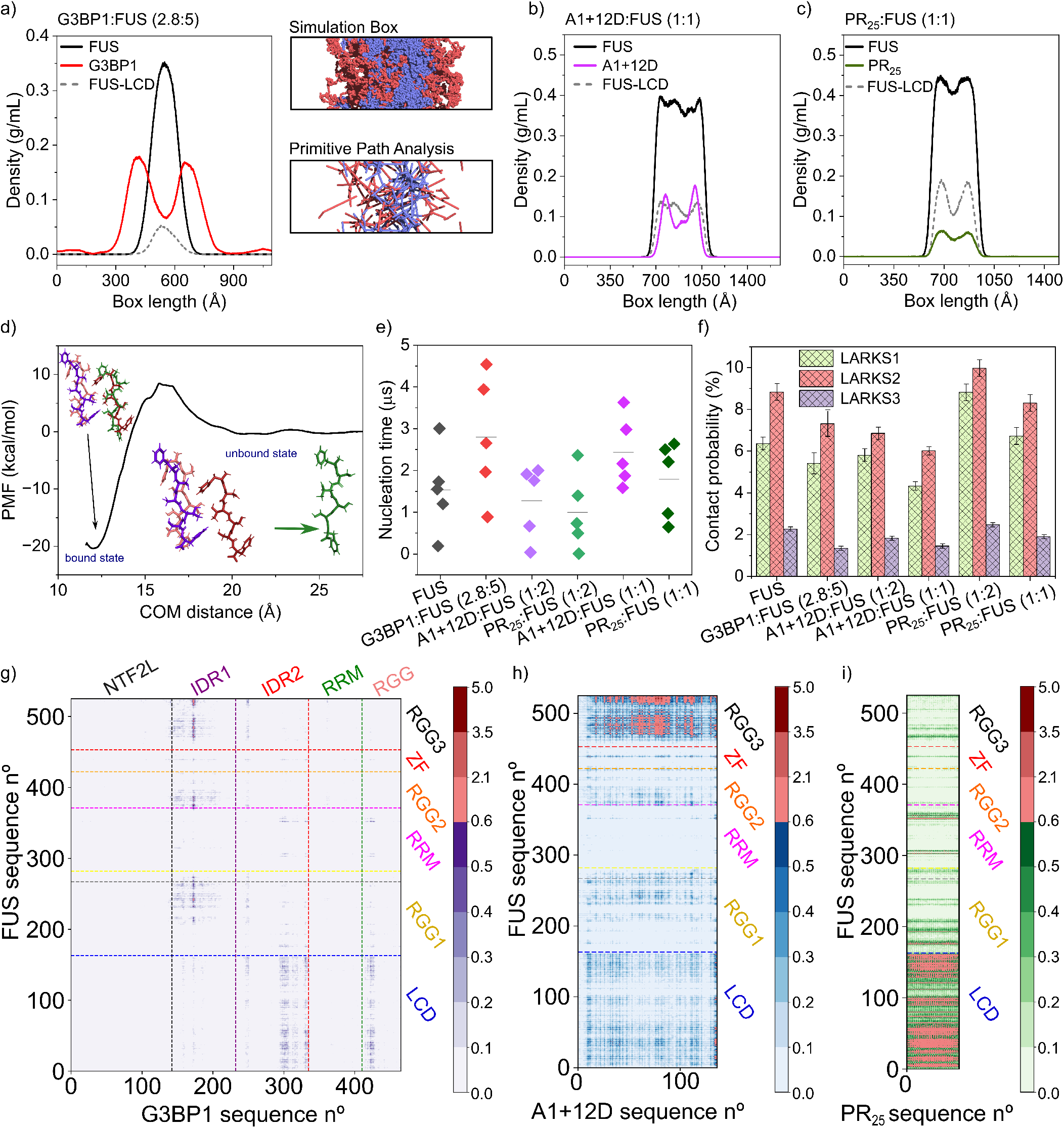
G3BP1 and highly charged sequences review FUS condensate organization and modulate their ageing kinetics. (a-c) Density profiles of G3BP1:FUS (a), A1+12D:FUS (b) and PR_25_:FUS (c) condensates at different molar ratios. Component distributions are shown for G3BP1 (red), PR_25_ (green), A1+12D (pink) and FUS, further decomposed into full-length (black) and LCD (grey) contributions. In (a), a snapshot of the DC simulation box and a primitive path analysis for a G3BP1:FUS mixture (2.8:5) are shown (G3BP1 in red and FUS in blue). (d) Potential of mean force (PMF) calculation as a function of the center-of-mass (COM) distance for the dissociation of the LARKS-forming G3BP1 peptide (^250^SWASVTSKNL^259^) from a cross-*β*-sheet stack composed of four peptides. (e) Average nucleation time (⟨*τ*⟩, black horizontal bar) for the formation of FUS-associated *β*-sheets in condensates containing FUS, G3BP1, A1+12D, and PR_25_ across different stoichiometries. Solid diamonds indicate nucleation times from independent trajectories where *β*-sheet transitions were observed. (f) Contact probability of intermolecular LARKS–LARKS interactions among FUS replicas in condensates formed by G3BP1:FUS, A1+12D:FUS, and PR_25_:FUS mixtures with different molar ratios. (g-i) Intermolecular contact maps of residue–residue contact frequencies (in %) of G3BP1-FUS (g) in a condensate with a 2.8:5 molar ratio, and of FUS–A1+12D (h) and FUS–PR_25_ (i), both under equimolar conditions. The FUS and G3BP1 domains are indicated in the maps.

An important question arising from the interfacial organization of G3BP1:FUS condensates is whether G3BP1 itself possesses an intrinsic propensity to undergo ageing through the formation of cross-*β*-sheet assemblies. Although G3BP1 has been implicated in stress granule dynamics, its ability to form amyloid-like structures remains controversial, and no well-characterized LARKS have been identified within its sequence^23,29,52^. To address this question, we systematically screened the IDRs of G3BP1 using ZipperDB^97^. This analysis identified the segment ^250^SWASVTSKNL^259^ as a strong candidate for steric-zipper formation. To evaluate the stability of this putative fibrillar motif, we performed atomistic MD simulations of a four-layer stack of the LARKS peptide using the a99SB-*disp*/TIP4P-*disp* force field^98^, following the strategy previously employed to characterize LARKS from other RNA-binding proteins^37,48,99^. We then computed the potential of mean force (PMF) associated with the dissociation of one peptide from the remaining cross-*β*-sheet stack as a function of the distance between their centers of mass (Section S4 of the SM). The resulting free energy profile (Fig. 3d) exhibits a binding free energy minimum of approximately − 20.4 kcal/mol, comparable to those previously reported for experimentally validated LARKS from FUS, NUP98, A*β*_42_ and other RNA-binding proteins such as hnRNPA1 or TDP43^37,38,48,99,100^. These results identify a previously unrecognized aggregation-prone motif within G3BP1, suggesting that this protein possesses an intrinsic molecular capacity to undergo progressive maturation.

We next investigated how the LARKS aggregation motifs in both G3BP1 and FUS influence the ageing of mixed condensates. To this end, we performed out-of-equilibrium DC simulations of G3BP1:FUS condensates using the dynamic ageing algorithm described above (Fig. 1d and Section S5 of the SM). Although the algorithm explicitly accounts for the LARKS of both proteins, no disorder-to-*β*-sheet transitions involving the G3BP1 LARKS were observed within the entire simulation timescale, indicating that the intrinsic aggregation propensity encoded within this sequence might not be sufficient to trigger structural hardening under near physiological conditions in comparable timescales to those for FUS. Fig. 3e shows the nucleation times of cross-*β* assemblies obtained from five independent trajectories of G3BP1:FUS condensates at a (2.8:5) molar ratio. Individual nucleation events (red diamonds) occur systematically later than in pure FUS condensates, resulting in an average delay of approximately 50% in cross-*β*- sheet formation. This stabilization of the liquid-like state emerges from the mesoscale architecture of the condensate. As shown in Fig. 3a, G3BP1 forms a persistent interfacial shell surrounding the FUS-rich phase while partially penetrating its interior, where it establishes heterotypic interactions with FUS. Recent work has shown that condensate interfaces constitute preferred nucleation hotspots for structural maturation^60,61,83^. Consequently, the G3BP1 coating does not simply alter condensate morphology; it selectively protects the most aggregation-prone regions of the condensed phase by reducing the accessibility of FUS molecules to these nucleation sites. At the same time, the partial penetration of G3BP1 into the FUS-rich core further disrupts homotypic FUS interactions, decreasing the local availability of LARKS contacts throughout the condensate. These results suggest that the reduced ageing of G3BP1:FUS condensates might not be explained solely by the intrinsic aggregation propensity encoded within the constituent protein sequences. Instead, they reveal a more general physical principle: ageing is governed by the interplay between sequence-encoded interaction motifs and the mesoscale organization of the condensed phase, which together determine whether aggregation-competent interaction networks emerge. In this framework, the interfacial organization generated by G3BP1 represents a distinct molecular strategy that delays structural hardening, by limiting the local availability of LARKS interactions required for cross-*β*-sheet nucleation.

We next investigated how highly charged sequences modulate the ageing of FUS condensates. Whereas G3BP1 decelerates ageing through interfacial organization, A1+12D and PR_25_ provide a complementary opportunity to isolate the contribution of electrostatic interactions within homogeneously mixed condensates. Fig. 3e summarizes the average nucleation times of cross-*β* assemblies for A1+12D:FUS and PR_25_:FUS condensates at different stoichiometries. We find that A1+12D exhibits a concentration-dependent protective effect. At a 1:2 molar ratio it produces only a marginal increase in the average nucleation time, whereas at a 1:1 ratio, maturation is delayed by approximately a two-fold. These results are fully consistent with the reduction in aggregation-prone LARKS contacts observed above (Fig. 2e), indicating that increasing concentrations of the negatively charged peptide progressively suppress the formation of aggregation-competent interaction networks. In contrast, PR_25_ produces the opposite behaviour at low concentration. At a 1:2 molar ratio, the peptide accelerates ageing by approximately 50%, whereas increasing its concentration to 1:1 restores nucleation times close to those of pure FUS condensates.

To understand the molecular origin of these distinct ageing behaviours, we quantified the probability of productive FUS LARKS–LARKS contacts in all multicomponent condensates (Fig. 3f). Remarkably, the trends in nucleation kinetics are faithfully reproduced by the local availability of aggregation-prone contacts. Whereas G3BP1:FUS and A1+12D:FUS condensates display reduced LARKS–LARKS contact probabilities relative to pure FUS, PR_25_:FUS condensates exhibit an increased probability of productive LARKS encounters, especially at the 1:2 molar ratio. Thus, despite acting through fundamentally different molecular mechanisms, the effects of RNA, hnRNPA1, G3BP1 and highly charged protein sequences all converge on the same microscopic descriptor of condensate ageing: the local availability of aggregation-competent interaction networks enriched in productive LARKS contacts.

To identify the molecular interactions responsible for these distinct behaviours, we next analyzed the intermolecular contact maps obtained from DC simulations of G3BP1:FUS, A1+12D:FUS and PR_25_:FUS condensates (Figs. 3g–i; Section S6 of the SM). Because G3BP1 is a homodimer composed of two identical 466-residue monomers, only the contacts corresponding to one monomer are shown in Fig. 3g. We find that the dominant heterotypic interactions occur between the FUS LCD and the IDR2 and RGG regions of G3BP1. This interaction pattern is readily explained by the sequence composition of these domains (Fig. S1 of the SM), as the arginine-rich IDR2 and RGG regions of G3BP1 establish favourable cation–*π* interactions with the aromatic residues of the FUS-LCD. In addition, the glutamate-rich IDR1 of G3BP1 interacts preferentially with the positively charged RGG domains of FUS through electrostatic attraction. Together, these interactions effectively compete with the intrinsic homotypic interactions that normally stabilize the FUS interaction network, thereby reducing the availability of productive LCD– LCD contacts required for disorder-to-order structural transitions. This competitive rewiring of the interaction network provides a molecular explanation for the protective role of G3BP1. Rather than simply coating the condensate, G3BP1 selectively redistributes the intermolecular interactions that govern FUS interprotein networks. Combined with the interfacial architecture described above, these heterotypic interactions partially preclude the emergence of aggregation-competent microenvironments by simultaneously shielding preferred nucleation sites and reducing the probability of LARKS encounters. Notably, this mechanism differs fundamentally from those identified for RNA and hnRNPA1, which primarily regulate condensate ageing through changes in condensate density and heterotypic mixing, respectively.

The intermolecular contact maps of condensates containing either A1+12D or PR_25_ also provide a molecular explanation for their distinct effects on FUS interprotein *β*-sheet formation. In the A1+12D:FUS (1:1) condensate (Fig. 3h), A1+12D interacts preferentially with the RGG3 domain of FUS, thereby competing with the intrinsic LCD–RGG interactions that stabilize the FUS homotypic interaction network. This competition weakens the alignment between FUS LCD and its RGG domains—which indirectly contributes to enhancing ageing as FUS LCD–LCD interactions become further available. However, at the same time it induces a high degree of heterotypic mixing (Fig. 3b) reducing the emergence of aggregation microenvironments enriched in productive LARKS contacts. Consequently, overall, cross-*β*-sheet nucleation is delayed. In contrast, at a 1:2 molar ratio, A1+12D is not sufficiently abundant to substantially rewire the intermolecular interaction network, explaining the minor changes observed in both nucleation kinetics (Fig. 3e) and LARKS contact probabilities (Fig. 2f). A distinct mechanism emerges for PR_25_. As shown in Fig. 3i, the peptide interacts predominantly with the FUS LCD through favourable cation–*π* interactions, coating the aggregation-prone regions of the protein. At low peptide concentrations (1:2), PR_25_ is sufficiently sparse to locally promote the alignment of neighbouring FUS LCDs without significantly competing for their interaction hotspots and precluding FUS LCD-RGG3 contacts. This local increase in LCD–LCD contacts explains the modest acceleration of structural transitions observed in Fig. 3e (light green diamonds). However, as the peptide concentration increases (1:1), PR_25_ progressively saturates the available LCD interaction sites, disrupting homotypic FUS LCD associations and reducing the formation of aggregation interaction networks. Consequently, the nucleation kinetics of cross-*β*-sheet formation returns to values comparable to those of pure FUS condensates. Together, these results demonstrate that highly charged sequences regulate condensate ageing through competition for key interaction hotspots within FUS. Whereas A1+12D slows down at increasing concentrations FUS ageing by rewiring the interaction network through the RGG domains, PR_25_ directly targets the LCD and exhibits a concentration-dependent dual behaviour, acting either as a transient promoter or as a inhibitor of aggregation depending on the condensate stoichiometry. These findings further reinforce the general idea emerging throughout this work, in which diverse molecular perturbations regulate condensate ageing by controlling the clustering of aggregation-competent interaction networks rather than simply altering the intrinsic aggregation propensity of the constituent proteins.

### D. RNA and G3BP1 cooperatively drive microphase separation within FUS condensates

G3BP1 is one of the principal scaffold proteins that nucleates stress granules by coordinating an extensive network of protein–protein and protein–RNA interactions^23,29,101^. Through its ability to bind RNA and recruit multiple RNA-binding proteins including FUS, G3BP1 plays a central role in defining both the composition and the internal organization of these condensates. Recent experimental studies have further demonstrated that RNA is not merely a passive client of stress granules, but an active regulator of their assembly, material properties, and mesoscale organization in a concentration-dependent manner^75,102,103^. Despite the central roles of G3BP1 and RNA in stress granule biology, it remains unknown how they cooperate to organize multicomponent condensates and regulate the microscopic interaction networks that govern pathological ageing. Here, we therefore investigate whether the combined action of G3BP1 and RNA gives rise to emergent condensate architectures beyond those observed for either component alone in presence of FUS, and how these architectures regulate the local availability of aggregation-prone contacts and the nucleation of cross-*β*-sheet structures.

First, we investigated how RNA concentration regulates the internal architecture of polyA:G3BP1:FUS condensates by analyzing density profiles obtained from DC simulations at two different polyA:G3BP1:FUS molar ratios: 1:(2.8):5 (Fig. 4a) and (2.6):(2.8):5 (Fig. 4b). Density profiles for G3BP1, polyA, and FUS are shown in red, teal green, and black, respectively, while the distribution of the FUS LCD is shown in grey (dashed curve). At the lower RNA concentration, polyA colocalizes mostly with the FUS-rich phase, whereas G3BP1 preferentially accumulates at the condensate interface, coating the FUS/polyA-rich core in a manner similar to that observed for binary G3BP1:FUS condensates (Fig. 3a). Interestingly, the FUS LCD exhibits partial exclusion from the RNA-rich region, indicating that RNA already reshapes the spatial distribution of aggregation-prone domains within the condensate. Increasing the RNA concentration produces a striking reorganization of the condensate architecture. Rather than simply expanding the RNA-rich phase, the system undergoes a microphase separation in which FUS and RNA segregate into distinct microdomains, while G3BP1 becomes homogeneously distributed throughout the condensate. This transition—evaluated through multiple independent trajectories simulated over 5 *mu*s each—is accompanied by a pronounced decrease in condensate density, consistent with the RNA-induced reentrant behaviour reported in Fig. 1b. Together, these results suggest that FUS, polyA and G3BP1 cooperate to generate emergent architectures that cannot be inferred from the behaviour of either component alone, revealing that condensate composition controls not only phase stability but also the spatial organization of their interaction networks.

**FIG. 4.**
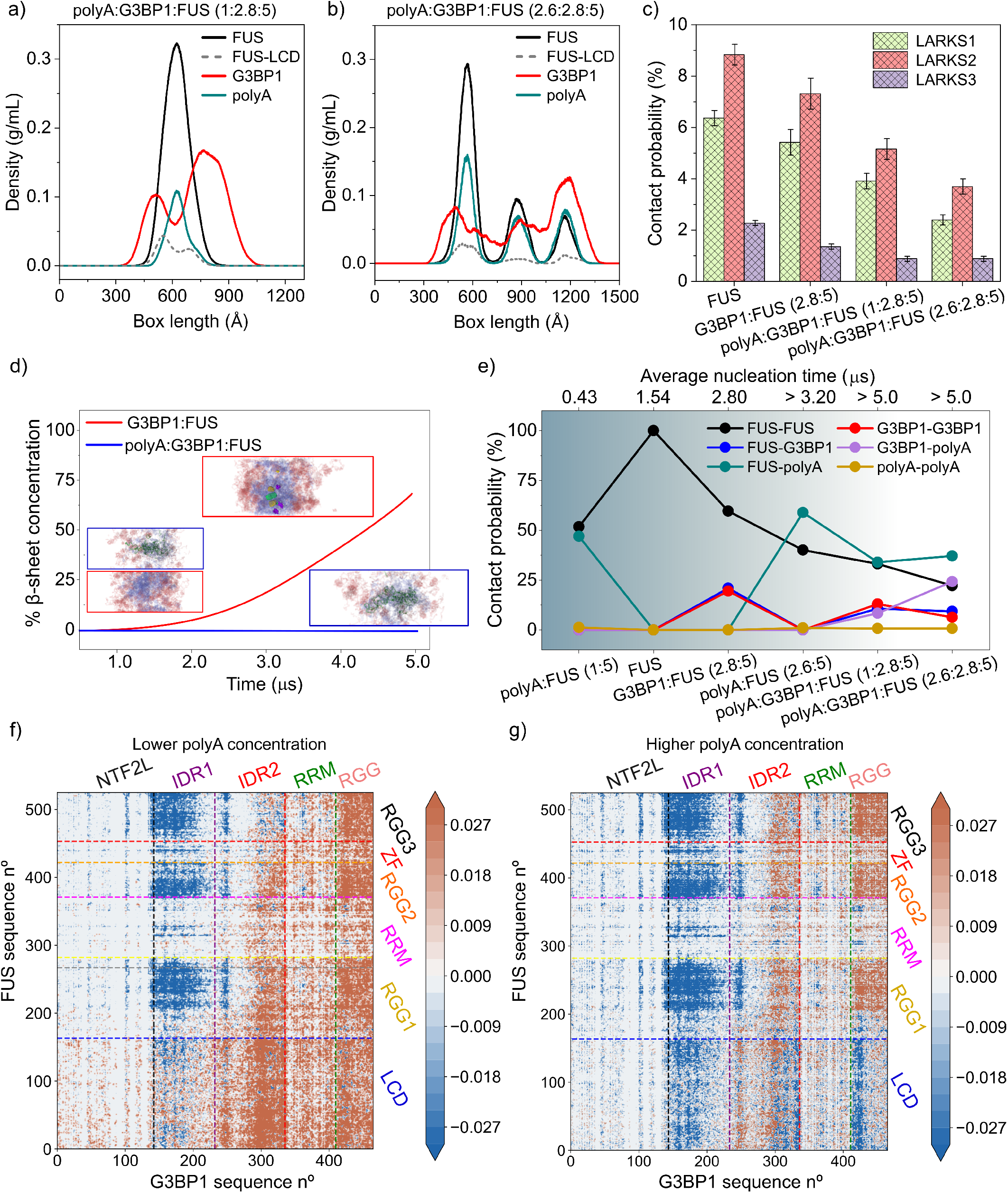
G3BP1/RNA-driven microphase-separation in FUS condensates prevents ageing. (a-b) Density profiles of polyA:G3BP1:FUS condensates with molar ratios of 1:2.8:5 (a) and 2.6:2.8:5 (b). Different component distributions for G3BP1 (red), polyA (teal), and FUS, separated into full-length (black) and LCD (grey) are depicted by different colours. (c) LARKS–LARKS contact probability of FUS molecules in condensates formed by FUS, G3BP1:FUS and polyA:G3BP1:FUS at different molar ratios. (d) Time-evolution of the cross-*β*-sheet concentration (including simulation snapshots) in G3BP1:FUS and polyA:G3BP1:FUS mixtures. (e) Total contact protein-protein probability (in %) of FUS, G3BP1, and polyA in condensates with different compositions and molar ratios (uncertainties are comparable to the marker size). (f-g) Intermolecular contact difference maps (in percentage variation) of FUS–G3BP1 calculated as the difference between a given polyA:G3BP1:FUS mixture ((f) at 1:2.8:5 molar ratio and (g) at 2.6:2.8:5 molar ratio) and the G3BP1:FUS reference contact map. Positive and negative differences are shown in orange and blue, respectively.

**FIG. 5.**
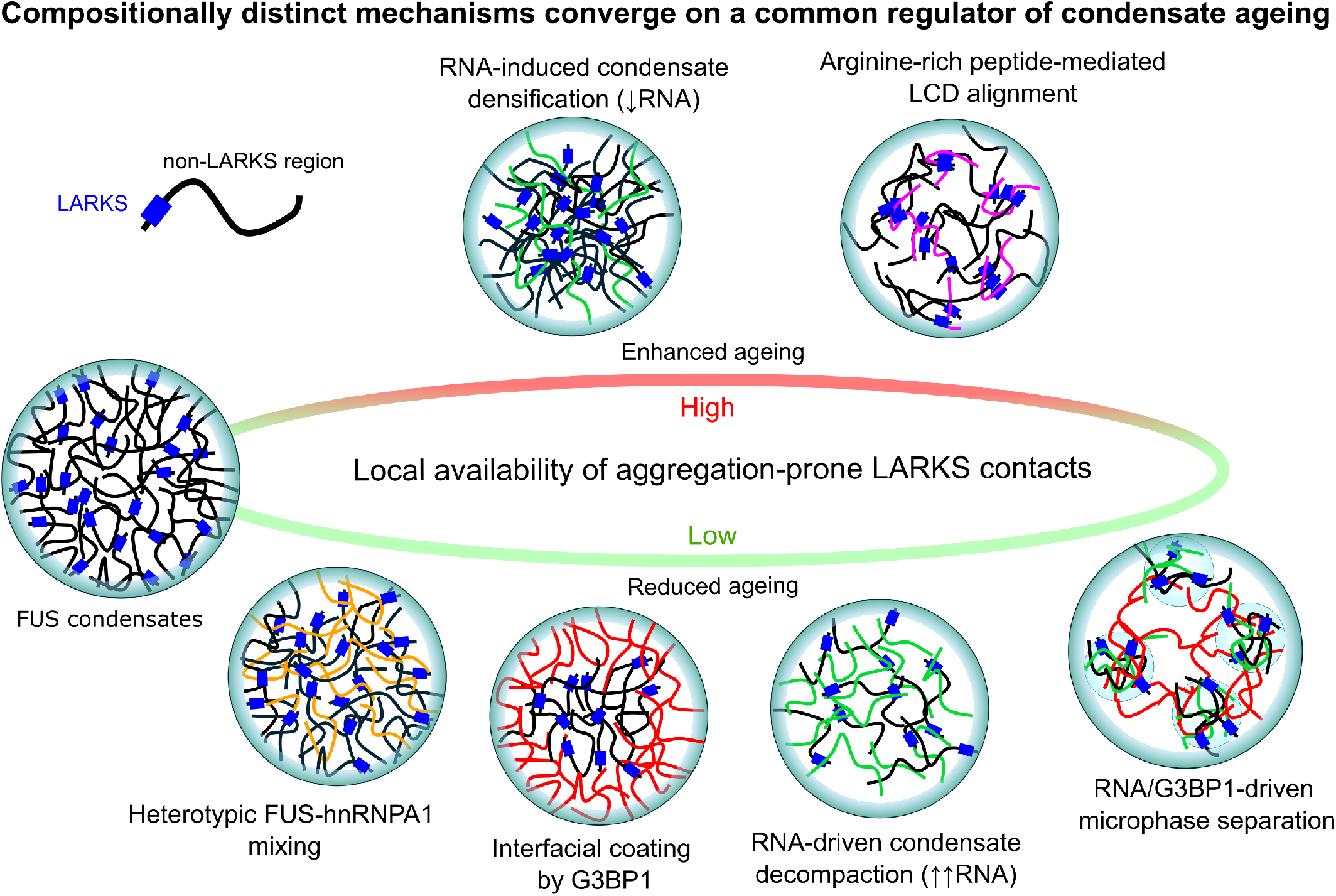
Molecular mechanisms governing cross-*β*-sheet formation in FUS-based multicomponent stress granules. Schematic summary of the molecular mechanisms by which condensate composition regulates ageing. Distinct compositional perturbations—including RNA concentration, heterotypic protein mixing, spatial organization, and G3BP1-mediated microphase separation—reconfigure the underlying intermolecular interaction network, thereby modulating the local availability of aggregation-prone LARKS contacts.

To determine how RNA further modulates ageing in G3BP1:FUS condensates, we quantified the probability of intermolecular FUS LARKS contacts from DC trajectories (Fig. 4c). Relative to binary G3BP1:FUS condensates, the incorporation of polyA leads to a pronounced reduction in the local availability of LARKS contacts, indicating that RNA further disrupts the microscopic network required for cross-*β*-sheet formation. To establish whether this molecular reorganization translates into enhanced resistance against kinetic arrest, we subsequently performed out-of-equilibrium simulations using the dynamic disorder-to-order transition algorithm. Five independent trajectories were simulated for each polyA:G3BP1:FUS composition (1:2.8:5 and 2.6:2.8:5). Remarkably, none of the 10 simulations exhibited formation of cross-*β* structures during the entire 5 *µ*s window of each simulation (Fig. 4d). This suppression of nucleation contrasts sharply with the behaviour of the corresponding binary systems and demonstrates that G3BP1 and RNA cooperatively delay the earliest molecular events leading to condensate ageing. Mechanistically, G3BP1 promotes extensive heterotypic interaction networks with both polyA and FUS, while polyA simultaneously reduces condensate density and reorganizes FUS intermolecular contacts. Together, these effects minimize the local availability of LARKS required for structural conversion. In that sense, our simulations suggest that physiological stress-granule components suppress pathological ageing not by eliminating aggregation-prone motifs, but by collectively rewiring condensate architecture to prevent the microscopic networks that initiate cross-*β* nucleation.

Although the density profiles and the analysis of FUS LARKS–LARKS interactions provide fundamental insight into the reduced propensity of RNA-containing condensates to undergo maturation, they do not directly quantify how the interaction network is redistributed among the different condensate components. As shown in the previous section, the degree of heterotypic mixing is itself a major determinant of ageing (Fig. 2). We therefore quantified the normalized global fraction of intermolecular contacts established by every pair of condensate components (Fig. 4e). This quantity provides a direct measure of condensate mixing, since a larger fraction of contacts between two components reflects a higher probability of their co-localization (see Section S6 of the SM for details on these calculations). The resulting interaction networks reveal a progressive rewiring of inter-molecular contacts as the RNA concentration increases. Most notably, homotypic FUS–FUS interactions decrease monotonically, whereas heterotypic G3BP1–polyA interactions become increasingly dominant.

Simultaneously, the relative abundance of FUS–polyA contacts decreases, indicating that RNA progressively exchanges FUS for G3BP1 as its preferred interaction partner. This redistribution explains the structural transition observed in the density profiles (Figs. 4a–b). At low RNA concentrations, G3BP1 retains a partially self-associated shell surrounding the FUS-rich core, whereas increasing the RNA content weakens G3BP1–G3BP1 contacts while strengthening G3BP1–polyA interactions. The resulting interaction network promotes condensate decompaction and the emergence of RNA- and FUS-rich microdomains, while G3BP1 acts as a molecular bridge that maintains condensate cohesion. From a mechanistic perspective, this interaction rearrangement has an important consequence. As G3BP1 and RNA progressively sequester intermolecular contacts away from FUS, the probability of forming persistent homotypic FUS interaction clusters decreases, thereby reducing the local availability of LARKS contacts required for cross-*β* nucleation.

After establishing that polyA hinders cross-*β*-sheet emergence in G3BP1:FUS condensates, we next sought to identify the molecular interactions responsible for this protective effect. To this end, we computed intermolecular FUS–G3BP1 contact maps and compared them with those of the corresponding RNA-free condensates. Rather than displaying the absolute contact maps, Figs. 4f–g show their differences relative to the G3BP1:FUS reference system (Fig. 3g), thereby directly highlighting the interactions that are selectively strengthened or weakened upon RNA addition. At low RNA concentration, one of the most prominent changes is the disappearance of interactions between the G3BP1 IDR1 and the positively charged RGG domains of FUS. This redistribution is readily explained by the preferential binding of the negatively charged polyA chains to the FUS RGG regions, in agreement with the FUS–polyA contact maps (Figs. S5–S6). Although these RNA-mediated interactions are insufficient to alter the overall condensate architecture, which remains as a single dense phase (Fig. 4a), they locally reorganize the spatial distribution of FUS domains. In particular, the association of RNA with the FUS RGG domains reduces the alignment of FUS LCDs with their own RGGs, and increases the local heterotypic interactions of FUS LCDs with the IDR2, RRM and RGG domains of G3BP1, thereby hindering cross-*β*-sheet formation.

A qualitatively different mechanism emerges at high RNA concentrations. Increased electrostatic repulsion between polyA strands drives condensate decompaction and promotes the formation of FUS- and RNA-rich microdomains (Fig. 4b). Under these conditions, G3BP1 no longer acts primarily as an interfacial coating protein but instead becomes distributed throughout the condensate, maintaining cohesion between the emerging microdomains. Consistent with this reorganization, interactions between the RGG domains of FUS and G3BP1 become dominant (Fig. 4g). Although interactions between positively charged regions might initially appear counterintuitive, they are naturally explained by the simultaneous binding of both RGG domains to polyA chains, which therefore act as multivalent molecular bridges connecting the two proteins. This interpretation is fully consistent with the polyA–G3BP1 and polyA–FUS contact maps shown in Figs. S4–S6, and explains how these relatively dilute condensates remain structurally cohesive despite their partial decompaction. Taken together, these results reveal that RNA does not simply weaken protein–protein interactions. Instead, it hierarchically redistributes the intermolecular interaction network, from residue-level electrostatic contacts to mesoscale condensate organization, ultimately controlling the local availability LARKS interactions. Such hierarchical coupling between molecular interactions, condensate architecture, and inter-protein *β*-sheet formation provides a unified physical framework for understanding how multicomponent condensates may resist pathological maturation.

## III. CONCLUSIONS

In this work, we combined residue-resolution coarse-grained molecular dynamics simulations with a nonequilibrium disorder-to-order ageing algorithm to investigate how the molecular composition of multicomponent ribonucleoprotein condensates regulates their progressive solidification. This framework enabled us to bridge molecular interactions, condensate organization, and the emergence of pathological cross-*β* structures across multiple classes of stress-granule-associated condensates. Our results revealed that composition regulates condensate ageing through a common microscopic principle despite acting through distinct physical mechanisms. RNA, heterotypic RNA-binding proteins, scaffold proteins, and charged peptides each reorganize condensate architecture in different ways, yet all ultimately depends on how they modulate the local availability of aggregation-prone domains involved in the nucleation of cross-*β* structures. Thus, rather than being determined by composition alone, the propensity of condensates to undergo ageing emerges from how composition rewires the intermolecular interaction network governing these transitions.

We first showed that RNA regulates both the thermodynamic stability and ageing of FUS condensates in a non-monotonic manner (Fig. 1). Intermediate RNA concentrations stabilize condensates through favorable protein–RNA interactions, increasing condensate density and consequently enhancing the frequency of LARKS contacts. In contrast, high RNA concentrations induce condensate decompaction through reentrant phase behaviour, reducing local LARKS density and delaying cross-*β* formation. Contact map analysis demonstrates that these effects originate from an RNA-driven reorganization of the interaction network, whereby preferential binding of RNAs to the FUS RGG domains redistributes the balance between FUS LCD–RGG and LCD– LCD interactions, ultimately controlling the accessibility of LARKS.

Our simulations further demonstrate that condensate stoichiometry provides an independent mechanism for regulating ageing (Fig. 2). Increasing the proportion of heterotypic protein partners, including hnRNPA1, A1+NLS, and A1+12D, progressively shifts the interaction network from homotypic to heterotypic contacts. This enhanced molecular mixing reduces persistent homotypic interactions between FUS LARKS and consequently hinders the nucleation of cross-*β* structures. These results establish that heterotypic mixing does not simply alter condensate composition but actively reshapes the connectivity landscape governing precise inter-protein domain interactions.

Beyond condensate composition and mixing, we also identify mesoscale spatial organization as an additional layer of regulation (Figs. 3 and 4). Although A1+12D and PR_25_ modify FUS condensate interaction networks, they generate fundamentally well-mixed internal architectures. In contrast, G3BP1 preferentially accumulates at the condensate interface, where it partially coats the FUS-rich phase and competes for interactions and sites with FUS through its intrinsically disordered regions. This organization substantially reduces the accessibility of FUS LARKS and delays ageing despite the intrinsic aggregation propensity predicted for G3BP1 itself through our all-atom simulations (Fig. 3d). Conversely, charged sequences such as A1+12D and PR_25_ regulate ageing by directly competing for or promoting interactions involving the FUS LCD, producing concentration-dependent inhibition or enhancement of LARKS contacts depending on their concentration. These results demonstrate that condensate architecture itself constitutes an active regulator of ageing rather than simply a structural consequence of phase separation.

Finally, our simulations of ternary polyA:G3BP1:FUS condensates revealed that RNA and G3BP1 cooperate to establish highly protective condensate organizations characteristic of stress granules (Fig. 4). Increasing RNA concentration progressively redirects intermolecular interactions toward G3BP1–RNA networks while simultaneously reducing persistent FUS–FUS contacts. At high RNA concentrations, this reorganization leads to condensate decompaction together with the emergence of RNA/FUS-rich clusters stabilized by G3BP1, completely suppressing cross-*β*-sheet nucleation over the timescales explored in our simulations. These observations suggest that stress-granule organization itself may represent an evolved mechanism to minimize the formation of pathological aggregation nuclei while preserving condensate integrity^104,105^.

Taken together, our simulations identified a unified molecular framework linking condensate composition, mesoscale organization and ageing. Across all perturbations investigated—including RNA concentration, protein stoichiometry, heterotypic mixing, scaffold-mediated interfacial organization, charged peptides, and RNA-rich stress-granule architectures—we find that distinct compositional changes converge on a single microscopic determinant: the local availability of aggregation-prone contacts. This common descriptor provides a mechanistic explanation for how chemically diverse condensates can exhibit ageing behaviour despite possessing fundamentally different interaction networks and internal organizations^11,52,106,107^. Beyond the specific systems investigated here, our work establishes a general computational strategy for connecting condensate composition with long-timescale structural transitions. More broadly, it provides a predictive framework for understanding how biomolecular condensates regulate their resistance to pathological solidification and offers quantitative principles for engineering condensate organizations to modulate ageing in neurodegenerative disease and other condensate-related pathologies.

## IV. ACKNOWLEDGEMENTS

The authors acknowledge Prof. Antonio Rey for his useful insights on this research work. E.P. acknowledges funding from European Social Fund Plus and the project PID2022-136919NA-C33 from the Spanish MICIU. A.F. acknowledges funding from the Ramon y Cajal fellowship (RYC2021-030937-I) and Spanish National Grant (PID2022-136919NA-C33). F. G. acknowledges funding from the MICIU MCIN/AEI/10.13039/501100011033, project references PID2022-136919NA-C33 and PID2025-169417NB-C21. P.L acknowledges funding from the European Union’s Horizon 2020 research and innovation program (grant agreement 101160499 to J. R. E). A. R. T. acknowledges funding from the European Union Horizon 2020 research and innovation programme (grant agreement 803326 to R.C.-G.) and from Ministerio de Ciencia e Innovacion under the Juan de la Cierva fellowship (JDC2024-053759-I). R.C.-G. acknowledges funding the UK Research Innovation (UKRI) Engineering and Physical Sciences Research Council (EPSRC) [EP/Z002028/1], following funding from the European Research Council (ERC) Consolidator Grant “ChromatinDroplets” under the European Union’s Horizon Europe research and innovation programme. J.R.E. also acknowledges funding from the Ramon y Cajal fellowship (RYC2021-030937-I), the Spanish National Agency for Research (PID2022-136919NA-C33 and PID2025-169417NB-C21), and the European Research Council (ERC) under the European Union’s Horizon Europe research and innovation program (grant agreement no. 101160499). J.R.E also acknowledges the CRIS Cancer Foundation for the research grant CRIS-CANCER-4332687.The authors acknowledge the computational resources provided by the Red Española de Supercomputación (RES) at the Barcelona Supercomputing Center (BSC), through projects FI-2025-3-0003, FI-2025-3-0065 and EHPC-REG-2025R02-173 on the MareNostrum5 supercomputer, and the computational resources at the CIEMAT Xula supercomputer through project FI-2026-1-0027.

## Notes

### Competing Interest Statement

The authors have declared no competing interest.

