## supporting information for "Compositional control of FUS condensate ageing through aggregation-prone interaction networks"

(Dated: July 31, 2026)

[*]

### S1. THE MPIPI-RECHARGED MODEL

The Mpipi-Recharged model is a residue-level coarse-grained force field for both protein and protein/RNA condensates [1]. Each amino acid or nucleotide is represented by a single bead connected to adjacent residues by harmonic bonds. Globular domains of the proteins are treated as rigid bodies whose beads are fixed at the $C_\alpha$ position from the corresponding Protein Data Bank (PDB). The potential energy is computed as the sum of pairwise bonded ($E_{\text{bonded}}$) and non-bonded ($E_{\text{non-bonded}}$) interactions as:

$$E = E_{\text{bonded}} + E_{\text{non-bonded}}. \tag{S1}$$

The intrinsically disordered regions (IDRs) are modeled as fully flexible polymers and these are connected to the globular domains. The bonded potential is written as:

$$E_{\text{bonded}}(r_{ij}) = \sum_{(ij)\in\mathcal{B}} k(r_{ij} - r_0)^2, \tag{S2}$$

where $\mathcal{B}$ denotes the set of bonded bead pairs, $r_{ij}$ is the distance between the connected beads, $r_0 = 3.81\,\text{Å}$ and $r_0 = 5.00\,\text{Å}$ are the equilibrium bond lengths for protein and RNA, respectively. The spring constant $k = 9.6\,\text{kcal}\cdot\text{mol}^{-1}\cdot\text{Å}^{-2}$.

Non-bonded interactions consist of the sum of the hydrophobic interaction and the electrostatic interaction. The hydrophobic interaction is given by the Wang–Frenkel (WF) potential [2] that accounts for short-ranged excluded-volume repulsion and long-ranged attraction. This potential is defined as:

$$E_{\text{WF}} = \sum_{i<j} u_{ij}^{\text{WF}}(r_{ij}), \tag{S3}$$

where the pair potential is defined as:

$$u_{ij}^{\text{WF}}(r_{ij}) = \begin{cases} \epsilon_{ij}\alpha_{ij}\left[\left(\dfrac{\sigma_{ij}}{r_{ij}}\right)^{2\mu_{ij}} - 1\right]\left[\left(\dfrac{R_{ij}}{r_{ij}}\right)^{2\mu_{ij}} - 1\right]^{2\nu_{ij}}, & r_{ij} < R_{ij}, \\ 0, & r_{ij} \geq R_{ij}, \end{cases} \tag{S4}$$

where

$$\alpha_{ij} = 2\nu_{ij}\left(\frac{R_{ij}}{\sigma_{ij}}\right)^{2\mu_{ij}}\left\{\frac{2\nu_{ij}+1}{2\nu_{ij}\left[\left(\frac{R_{ij}}{\sigma_{ij}}\right)^{2\mu_{ij}} - 1\right]}\right\}^{2\nu_{ij}+1}. \tag{S5}$$

Here $\sigma_{ij}$ is the pair-of-beads diameter, obtained from the individual bead diameters using the Lorentz arithmetic mixing rule (i.e., $\sigma_{ij} = (\sigma_i + \sigma_j)/2$). $R_{ij} = 3\sigma_{ij}$ is the cut-off distance for the $ij$-th interaction. The interaction parameter $\epsilon_{ij}$ is defined for each specific amino acid pair based on our atomistic Potential of Mean Force calculations and bioinformatics data [1]. The exponent $\nu_{ij}$ is set to 1 for all pairs and $\mu_{ij}$ depends on the specific pair, ranging from 2 to 12 (see Ref. [1]). Notice that higher values of $\mu_{ij}$ lead to a steeper increase in the repulsive part of the potential. The interaction involving globular domains are reduced to account for the 'buried' interactions. In particular, the interaction between flexible regions and globular domains are screened by a factor of $\sqrt{0.7}$ and the WF interaction between residues in globular regions is scaled down a factor of 0.7.

The electrostatic contribution to the non-bonded interactions is described by a Yukawa (Y) potential. The total electrostatic interaction energy is given by:

$$E_{\text{electrostatic}} = \sum_{i<j} u_{ij}^{\text{Y}}(r_{ij}), \tag{S6}$$

where the pair potential is defined as:

$$u_{ij}^{\text{Y}}(r_{ij}) = \begin{cases} \dfrac{A_{ij}}{r_{ij}} \exp(-\kappa r_{ij}), & r_{ij} < r_c, \\ 0, & r_{ij} \geq r_c. \end{cases} \tag{S7}$$

Here, $A_{ij}$ is the pair-specific electrostatic interaction parameter and $\kappa$ is the inverse Debye screening length. The cutoff distance for the electrostatic interaction is set to $r_c =$ 3.5 nm. For a monovalent electrolyte, the inverse Debye screening length is expressed as $\kappa = \sqrt{8\pi\ell_B c_s}$, where $c_s$ is the salt concentration and $\ell_B = \frac{e_0^2}{4\pi\varepsilon_0\varepsilon_r k_B T}$ is the Bjerrum length. Unless otherwise stated, the salt concentration is set to 150 mM. The relative dielectric constant $\varepsilon_r$ varies with temperature according to the empiric formula [3]:

$$\varepsilon_r(T) = \frac{5321}{T} + 233.760 - 0.9297T + 1.417 \cdot 10^{-3}T^2 - 8.292 \cdot 10^{-7}T^3, \tag{S8}$$

for $T$ in Kelvin. The pair-specific optimized values of $A_{ij}$ of the Yukawa potential can be found in Ref. [1]. In the Mpipi-Recharged model, such parameters indicate that the interaction between oppositely charged pairs is significantly stronger than those between identically charged pairs.

### S2. PROTEIN SEQUENCES AND AMINO ACID DISTRIBUTION PATTERNS

In this section, the regions corresponding to the considered low-complexity aromatic-rich kinked segments (LARKS) are highlighted in blue for each protein sequence.

**FUS**

MASNDYTQQATQSYGAYPTQPGQGYSQQSSQPYGQQSYSGYSQSTDTSGYGQSSYSSYGQSQNTG
YGTQSTPQGYGSTGGYGSSQSSQSSYGQQSSYPGYGQQPAPSSTSGSYGSSSQSSSYGQPQSGSYSQ
QPSYGGQQQSYGQQQSYNPPQGYGQQNQYNSSSGGGGGGGGGNYGQDQSSMSSGGGSGGGYG
NQDQSGGGGSGGYGQQDRGGRGRGGSGGGGGGGGGGYNRSSGGYEPRGRGGGRGGRGGMGGS
DRGGFNKFGGPRDQGSRHDSEQDNSDNNTIFVQGLGENVTIESVADYFKQIGIIKTNKKTGQPMIN
LYTDRETGKLKGEATVSFDDPPSAKAAIDWFDGKEFSGNPIKVSFATRRADFNRGGGNGRGGRGR
GGPMGRGGYGGGGSGGGGRGGFPSGGGGGGGQQRAGDWKCPNPTCENMNFSWRNECNQCKA
PKPDGPGGGPGGSHMGGNYGDDRRGGRGGYDRGGYRGRGGDRGGFRGGRGGGDRGGFGPGK
MDSRGEHRQDRRERPY

We have used two PDB codes to simulate the globular regions of FUS: residues from 285-371 (PDB code: 2LCW) and from 422-453 (PDB code: 6G99).

**G3BP1**

MVMEKPSPLLVGREFVRQYYTLLNQAPDMLHRFYGKNSSYVHGGLDSNGKPADAVYGQKEIHRK
VMSQNFTNCHTKIRHVDAHATLNDGVVVQVMGLLSNNNQALRRFMQTFVLAPEGSVANKFYVHN
DIFRYQDEVFGGFVTEPQEESEEEVEEPEERQQTPEVVPDDSGTFYDQAVVSNDMEEHLEEPVAE
PEPDPEPEPEQEPVSEIQEEKPEPVLEETAPEDAQKSSSPAPADIAQTVQEDLRTFSWASVTSKNLP
PSGAVPVTGIPPHVVKVPASQPRPESKPESQIPPQRPQRDQRVREQRINIPPQRGPRPIREAGEQGD
IEPRRMVRHPDSHQLFIGNLPHEVDKSELKDFFQSYGNVVELRINSGGKLPNFGFVVFDDSEPVQK
VLSNRPIMFRGEVRLNVEEKKTRAAREGDRRDNRLRGPGGPRGGLGGGMRGPPRGGMVQKPGF
GVGRGLAPRQMVMEKPSPLLVGREFVRQYYTLLNQAPDMLHRFYGKNSSYVHGGLDSNGKPAD
AVYGQKEIHRKVMSQNFTNCHTKIRHVDAHATLNDGVVVQVMGLLSNNNQALRRFMQTFVLAPE
GSVANKFYVHNDIFRYQDEVFGGFVTEPQEESEEEVEEPEERQQTPEVVPDDSGTFYDQAVVSND
MEEHLEEPVAEPEPDPEPEPEQEPVSEIQEEKPEPVLEETAPEDAQKSSSPAPADIAQTVQEDLRTF
SWASVTSKNLPPSGAVPVTGIPPHVVKVPASQPRPESKPESQIPPQRPQRDQRVREQRINIPPQRGP
RPIREAGEQGDIEPRRMVRHPDSHQLFIGNLPHEVDKSELKDFFQSYGNVVELRINSGGKLPNFGFV

VFDDSEPVQKVLSNRPIMFRGEVRLNVEEKKTRAAREGDRRDNRLRGPGGPRGGLGGGMRGPPR
GGMVQKPGFGVGRGLAPRQ

We have used one PDB code to simulate the globular region of the dimerization domain of G3BP1: residues from 7-41, 52-117, 124-137, 473-507, 518-583, 590-603 (PDB code: 3Q90), and we have used AlphaFold for their RRM domains: residues from 338-413 and 804-879 with a confidence level of 90%.

**hnRNPA1**

MSKSESPKEPEQLRKLFIGGLSFETTDESLRSHFEQWGTLTDCVVMRDPNTKRSRGFGFVTYATV
EEVDAAMNARPHKVDGRVVEPKRAVSREDSQRPGAHLTVKKIFVGGIKEDTEEHHLRDYFEQYG
KIEVIEIMTDRGSGKKRGFAFVTFDDHDSVDKIVIQKYHTVNGHNCEVRKALSKQEM ASASSSQR
GRSGSGNFGGGRGGGFGGNDNFGRGGNFSGRGGFGGSRGGGGYGGSGDGYNGFGNDGGYGGG
GPGYSGGSRGYGSGGQGYGNQGSGYGGSGSYDSYNNGGGGGFGGGSGSNFGGGGSYNDFGNYN
NQSSNFGPMKGGNFGGRSSGPYGGGGQYFAKPRNQGGYGGSSSSSSYGSGRRF

We have used the PDB code to simulate two globular regions of hnRNPA1, residues from 9–91 and from 103–181, both in the same PDB (PDB code: 1L3K).

**A1+NLS**

GSM ASASSSQRGRSGSGNFGGGRGGGFGGNDNFGRGGNFSGRGGFGGSRGGGGYGGSGD GYN-
GFGNDGGYGGGGPGYSGGSRGYGSGGQGYGNQGSGYGGSGSYDSYNNGGGGGFGGGSGSNFGG
GGSYNDFGNYNNQSSNFGPMKGGNFGGRSSGPYGGGGQYFAKPRNQGGYGGSSSSSSYGSGRRF

**A1+12D**

GSMASADSSQRDRDDSGNFGDGRGGGFGGNDNFGRGGNFSDRGGFGGSRGDGGYGGDGDGYNG
FGNDGSNFGGGGSYNDFGNYNNQSSNFDPMKGGNFGDRSSGPYDGGGQYFAKPRNQGGYGGSSS
SSSYGSDRRF

**polyA**

A 200-nucleotide polyadenine homopolymeric sequence.

### $PR_{25}$

Sequence composed of 25 repeats of the proline-arginine dipeptide (PR).

#### Amino acid distribution patterns

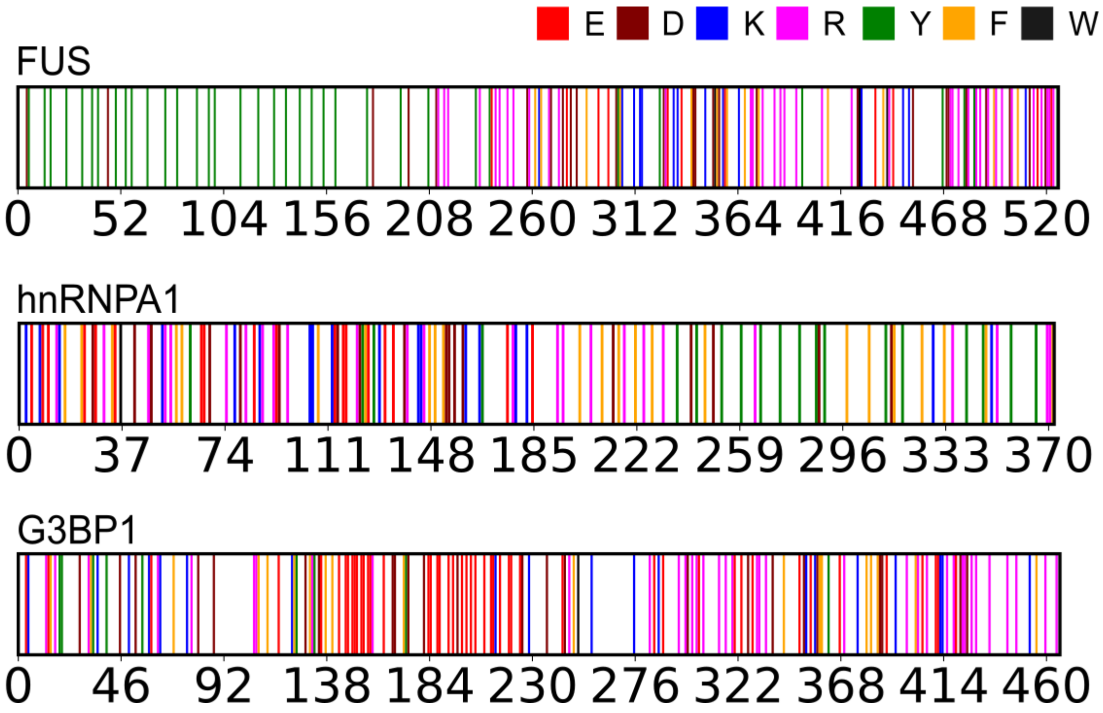


**FIG. S1:** Schematic description of the charged and aromatic residues along the sequences of FUS, hnRNPA1, and G3BP1. Glutamic acid (E) residues are shown in red, aspartic acid residues (D) in maroon, lysine residues (K) in blue, arginine residues (R) in pink, tyrosine residues (Y) in green, phenylalanine residues (F) in yellow, and tryptophan residues (W) in black.

### S3. DIRECT COEXISTENCE SIMULATIONS, DENSITY PROFILES AND PHASE DIAGRAMS

All direct coexistence (DC) simulations [4, 5] were performed following the protocol described below. These simulations were used to compute density profiles, phase diagrams, and other observables analysed throughout this work. The proteins were placed in a prismatic elongated box to simulate both the high-density and low-density phases separated by an interface. The long side of the box is perpendicular to the interfaces. Since determination of the optimal box dimensions is key to minimising finite size effects [6] while ensuring computer efficiency, we provide the following guidelines:

1. We choose the number of molecules of each component depending on the specific protein system to ensure stable phase coexistence. For example, for the pure FUS condensates, we use 49 FUS molecules, whereas for the polyA:FUS (1:5) condensates, we use 10 polyA chains of 200 nucleotides each and 49 FUS molecules.

2. We enforce the short sides of the slab box to be at least twice the radius of gyration of the proteins in the system to avoid self-interactions across the periodic boundary conditions. For example, for the pure FUS condensates, we use a slab box with a cross section of $15.4 \times 15.4$ nm$^2$ and 173.2 nm along the long side and for the polyA:FUS system we use a box with a cross section of $20.3 \times 20.3$ nm$^2$ and 130.0 nm along the long side.

3. The long side of the slab box should keep the total density of the system at approximately $\sim 0.1\,\mathrm{g{\cdot}cm^{-3}}$.

Simulations are carried out using the molecular dynamics package LAMMPS [7] (version 2 August 2023). We perform simulations in the canonical ($NVT$) ensemble using a Nosé-Hoover thermostat [8] for the rigid bodies (representing the globular domains) included in the RIGID package, and a Langevin thermostat [9] for the particles in flexible regions, both with a relaxation time of 5 ps. The time step for the Verlet integration of the equations of motion is 10 fs. After an equilibration period ($\sim 1 - 1.5$ $\mu$s), production runs of $1 - 2$ $\mu$s are performed depending on the specific system.

The phase diagrams of the systems was characterised from the density profiles obtained from the DC simulations. Density profiles were computed by averaging the mass density along the long axis of the simulation box, i.e., the direction normal to the slab interfaces. When two coexisting phases are present, the densities of the high-density and low-density phases are extracted from the corresponding plateau regions of these density profiles. These values were subsequently used to construct the phase diagrams reported throughout this work. Moreover, the critical density ($\rho_c$) and temperature ($T_c$) of the phase diagrams are evaluated by means of the law of rectilinear diameters and critical exponents [10, 11]:

$$\frac{\rho_l(T) + \rho_d(T)}{2} = \rho_c + s_2(T_c - T), \tag{S9}$$

and

$$(\rho_l(T) - \rho_d(T))^\beta = s_1\left(1 - \frac{T}{T_c}\right), \tag{S10}$$

where the critical exponent $\beta = 3.06$ for the three-dimensional Ising model [11], $\rho_d$ and $\rho_l$ are the coexisting densities of the dilute and condensed phases, respectively, and $s_1$ and $s_2$ are fitting parameters.

The $\rho_c$ and $T_c$ are estimated from DC simulations conducted at different temperatures, spaced by 10 K intervals. At each temperature, the density profile is analyzed to identify whether phase separation occurs. Thus, when two consecutive simulations are observed—one showing phase separation and the other not—it is inferred that the critical temperature lies within the intermediate range, with an uncertainty of approximately ±5 K. This approximation is reasonable, as the numerical error introduced by the numeric methods is negligible compared to the temperature steps used.

### S4. POTENTIAL OF MEAN FORCE CALCULATIONS

A LARKS-like candidate segment in the G3BP1 intrinsically disordered region (IDR) was identified using ZipperDB [12]. The predicted sequence was $^{250}$SWASVTSKNL$^{259}$ and exhibited steric zipper propensity with an interface energy below $-23$ kcal·mol$^{-1}$. Fibrillar assemblies were constructed for each identified motif using PyRosetta [13] in a cross-$\beta$ configuration. Fibrillar arrays consisting of 10 G3BP1 chains in cross-$\beta$ steric zipper configuration, percolated along the simulation box principal axis, were pre-equilibrated for 3 ns at 300 K. Four central chains from the equilibrated fibrillar array were subsequently selected for umbrella sampling potential of mean force (PMF) calculations.

Molecular dynamics simulations were performed with the Amber99sb-disp force field [14] in GROMACS 2023 [15]. Systems were solvated in orthorhombic boxes ($5 \times 5 \times 12$ nm) at 150 mM NaCl, with ion parameters validated for aqueous solubility at 300 K. After energy minimization (maximum force $< 1000$ kJ·mol$^{-1}$·nm$^{-1}$), hydrogen bonds were constrained using LINCS [16] with a 2 fs timestep. Electrostatics were treated with Particle-Mesh Ewald (PME) [17] (real-space cutoff 0.9 nm) under periodic boundary conditions. Temperature

(300 K, velocity-rescale thermostat, $\tau_T = 1$ ps) and pressure (1 atm, Parrinello–Rahman barostat, $\tau_P = 1$ ps) were held constant.

Potentials of mean force for G3BP1 fibril were computed via umbrella sampling by pulling a single chain away from the fibrillar core along the separation coordinate (distance from core), with the remaining three chains held in place. Center-of-mass (COM) distance was biased using a harmonic umbrella potential with a force constant of 10000 kJ mol$^{-1}$ nm$^{-2}$. Approximately 40 umbrella windows were generated at 0.025 nm intervals along the reaction coordinate. During production runs, positional restraints of 1000 kJ mol$^{-1}$ nm$^{-2}$ were applied (perpendicular to the pulling axis) to heavy atoms of selected protein chains to prevent overall rotation. Relative dissociation free-energy profiles along a constrained separation coordinate were reconstructed using WHAM [18]. A total accumulated simulation time of 500 ns was employed.

### S5. DYNAMIC ALGORITHM FOR CONDENSATE AGEING SIMULATIONS

We perform our aging simulations allowing the formation of inter-protein $\beta$-sheets to take place, according to the scheme developed in Refs. [19, 20] We first identified the regions of the proteins sequence that are prone to forming inter-protein secondary structures, known as LARKS. In the FUS sequence we consider the following LARKS [21]: $^{37}$SYSGYS$^{42}$, $^{54}$SYSSYGQS$^{61}$), and $^{77}$STGGYG$^{82}$, while for the G3BP1 sequence we consider the LARKS $^{250}$SWASVTSKNL$^{259}$ as described in the section S4. To enable "effective" disorder-to-order transitions by forming inter-protein $\beta$-sheets between the LARKS of the proteins the algorithm checks the local environments of the LARKS every $10^2$ simulation time steps. When three LARKS within the coordination cut-off distance ($r_{cut}$, see Table S1), the dynamic algorithm changes the force field parameters ($\epsilon_{ij}$,$\sigma_{ij}$) of the bead contained in the LARKS to those ($\epsilon_{ij,ordered}$,$\sigma_{ij,ordered}$) given in Table S1. The parameters correspond to inter-protein structured $\beta$-sheets as obtained from PMF calculations of atomistic simulations (see Refs. [19, 20, 22] and section S4). Although these all-atom simulations [19] showed that a minimum of four LARKS is required to form a stable fibrillar $\beta$-sheet unit, its strict implementation is overly restrictive over the timescales accessible for full-length proteins [23]. For this reason, we adopted a reduced value of three LARKS per event, which improves the sampling of nucleation events without modifying the underlying LARKS-mediated aggregation mechanism. The formed inter-protein $\beta$-sheet can grow if another LARKS is recruited within the $r_{cut,cross}$ distance (see Table S1). Similarly, an inter-protein $\beta$-sheet is reverted if one of the structured LARKS separates from the crossed-$\beta$-sheet motif a distance ($r_{cut,rev}$ away, see Table S1). Additionally, the algorithm increases the local stiffness to mimic the structured $\beta$-sheet by introducing a harmonic angular potential given by:

$$E_{\text{angle}} = \sum_{\text{angles}} k_{ang}(\theta - \theta_0)^2, \tag{S11}$$

where we set $\theta_0 = 180°$ and $k_{ang}$=5 kcal mol$^{-1}$ rad$^{-2}$ for the structured LARKS of an inter-protein $\beta$-sheet.

| | LARKS of FUS | | | LARKS of G3BP1 |
|---|---|---|---|---|
| | $^{37}$SYSGYS$^{42}$ | $^{54}$SYSSYGQS$^{61}$ | $^{77}$STGGYG$^{82}$ | $^{250}$SWASVTSKNL$^{259}$ |
| Central amino acid | $S_{39}$ | $S_{57}$ | $G_{79}$ | $S_{253}$ / $S_{719}$ |
| $m_{ordered}$ / (g mol$^{-1}$) | 107.44 | 107.48 | 86.92 | 107.42 |
| $\epsilon_{ij,ordered}$ / (kcal mol$^{-1}$) | 1.551 | 3.113 | 0.892 | 1.551 |
| $\sigma_{ij,ordered}$ / Å | 5.733 | 5.761 | 5.353 | 5.986 |
| $r_{cut}$ / Å | 11.20 | 11.20 | 11.20 | 11.20 |
| $r_{cut,cross}$ / Å | 11.50 | 11.50 | 11.50 | 11.50 |
| $r_{cut,rev}$ / Å | - | - | 20 | - |
| Coordination number | 3 | 3 | 3 | 3 |

**TABLE S1:** Parameters employed for residues belonging to structured inter-peptide $\beta$-sheet motifs in Mpipi-Recharged simulations for the LARKS in FUS and G3BP1, including the mass $m_{ordered}$, the interaction $\epsilon_{ij,ordered}$ and the steric radius $\sigma_{ij,ordered}$ of the structured LARKS, the cut off distances ($r_{cut}$, $r_{cut,cross}$, $r_{cut,rev}$), and the number of LARKS necessary to trigger the algorithm (coordination number).

These simulation were performed using the bond/react fix available in the REACTION package [24] of LAMMPS (version 2 Aug 2023), which allows us to change the topology and identity of the selected residues (i.e., LARKS within a LARKS high-density fluctuation) in a time- and local-dependent way. We run the dynamic algorithm in the isothermal-isobaric ($NpT$) ensemble for polyA:FUS systems with a relaxation time of 5 ps for the Langevin thermostat and the Nosé Hoover barostat, and a time step of 10 fs. Moreover, we perform these simulations in the $NVT$ ensemble under direct coexistence conditions for all other systems, using a relaxation time of 5 ps for the Langevin thermostat and a time step of 10 fs.

### S6. CALCULATION AND ANALYSIS OF CONTACT MAPS

Intermolecular contact maps within protein condensates were calculated from 1-3 $\mu$s $NVT$ trajectories at 290 K and at a monovalent salt concentration of 150 mM. The contact frequency between each pair of amino acids was calculated by summing the number of intermolecular contacts observed over all frames of the trajectories and normalizing by the total number of frames and the number of proteins in the system. Typically, molecular contacts are identified based on a distance criterion, with the assumption that the relative frequency of contact map occurrences (rather than absolute frequency) remains generally unaffected by the selected cut-off distance used in calculations, provided the cut-off values are reasonable. However, to accurately capture the most relevant and common residue-residue contact pairs that facilitate the phase separation, it is highly recommended to account for the specific parameterisation of each amino acid in terms of excluded volume and minimum potential energy interaction distance. Hence, we adopted a sequence-dependent cut-off distance equivalent to $1.2\sigma_{ij}$, where $\sigma_{ij}$ represents the mean excluded volume of the respective $i$-th and $j$-th amino acid [25]. Given the short-range nature of the Wang-Frenkel interactions in the Mpipi-Recharged model, the minima of the corresponding pair-specific potentials are located close to $2^{1/6}\sigma_{ij}$ ($\approx 1.122\sigma_{ij}$), with slight variations depending on the amino acid pair. Therefore, the cut-off distance is set to $1.2\sigma_{ij}$ to ensure that the relevant attractive interactions are captured. By implementing this sequence-dependent cut-off scheme for each amino acid pair interaction, we can reduce the contribution of non-specific short-range contacts, thus enhancing our ability to accurately identify the amino acids that positively contribute to condensate formation [25].

The resulting intermolecular contact maps show the total contact frequency between the amino acids of different proteins within a condensate. To quantify the protein–protein or LARKS–LARKS interactions, we sum the elements of the contact map corresponding to the amino acid pairs of interest. For protein-protein interactions, we consider all amino acid pairs between different proteins, whereas for LARKS–LARKS interactions, we restrict the analysis to the amino acids belonging to the identified LARKS regions.

### S7. ANALYSIS OF TOPOLOGICAL CONSTRAINTS USING PRIMITIVE PATH ANALYSIS

We analyze the network connectivity of the condensates using the topological constraints identified by Primitive Path Analysis (PPA). We use an equilibrated configuration extracted from an $NVT$ trajectory of the system as the input structure for the PPA analysis, which is performed using the PPA method [26], modified in Tejedor *et al.* [20]. The PPA algorithm minimizes the contour length of all chains in the condensate while keeping the terminal residues of the monomers fixed and, in this way, preserving the network topology by preventing chains from crossing each other. The energy minimization is performed with a tolerance of $10^{-7}$ for both the energy and the force, allowing up to $2 \times 10^5$ iterations. The resulting primitive paths identify the topological constraints between proteins, which we use to define the connectivity network of the condensate and characterize its structural organization.

### S8. SUPPORTING FIGURES: DENSITY PROFILES AND CONTACT MAPS

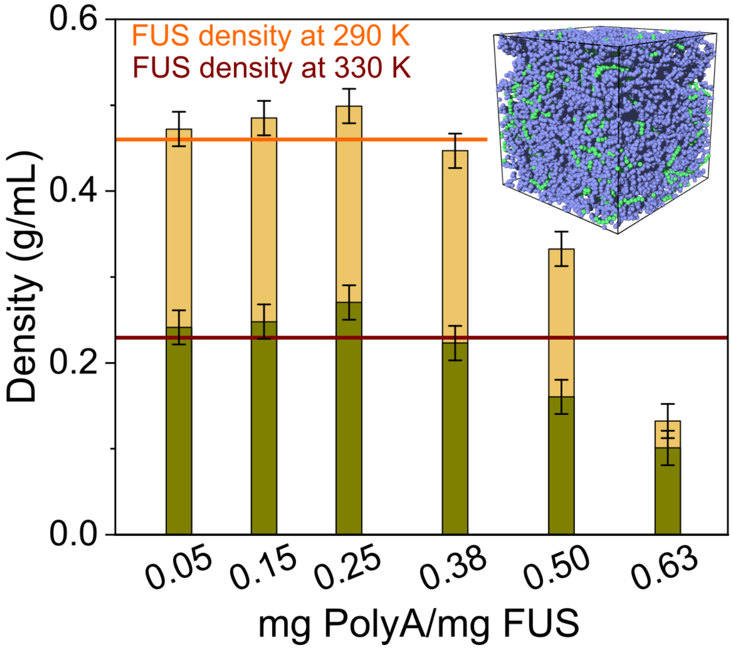


**FIG. S2:** Density of polyA:FUS mixtures as a function of the RNA/FUS mass ratio evaluated at 290 and 330 K in orange and olive green, respectively.

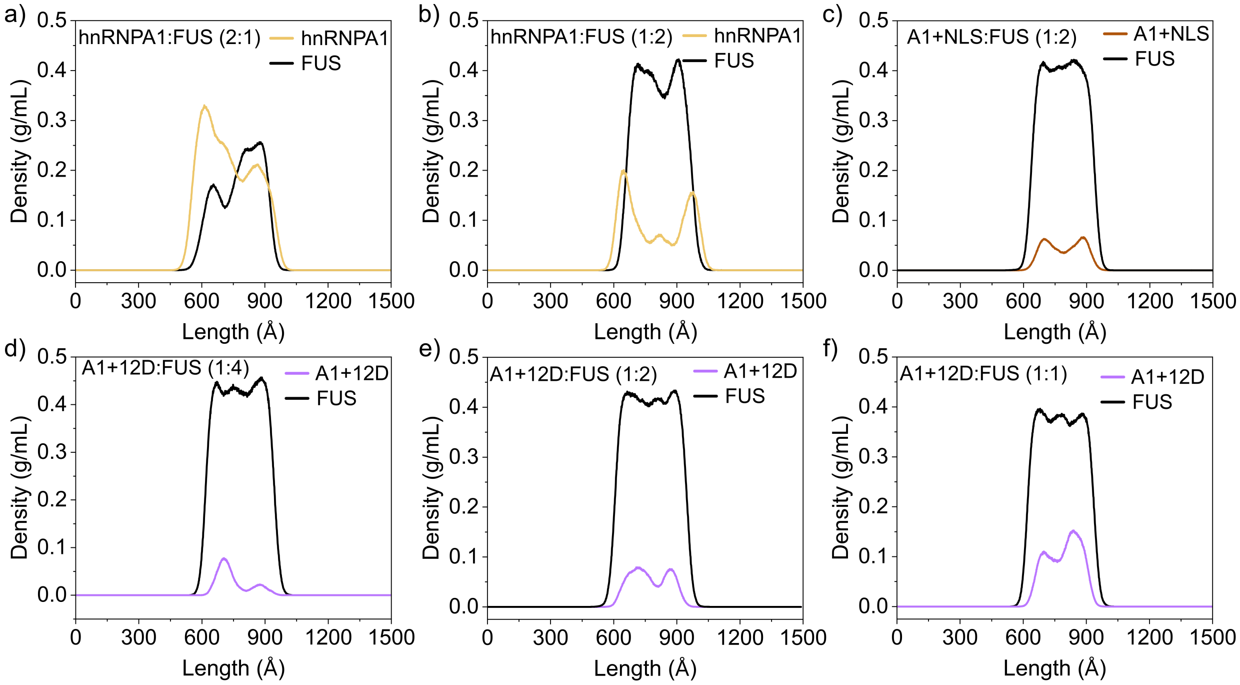


**FIG. S3:** Density profiles of condensates composed of hnRNPA1:FUS (2:1) in (a), hnRNPA1:FUS (1:2) in (b), A1+NLS:FUS (1:2) in (c), A1+12D:FUS (1:4) in (d), A1+12D:FUS (1:2) in (e) and A1+12D:FUS (1:1) in (f). The $x$ axis represents the length of the direct coexistence simulation box.

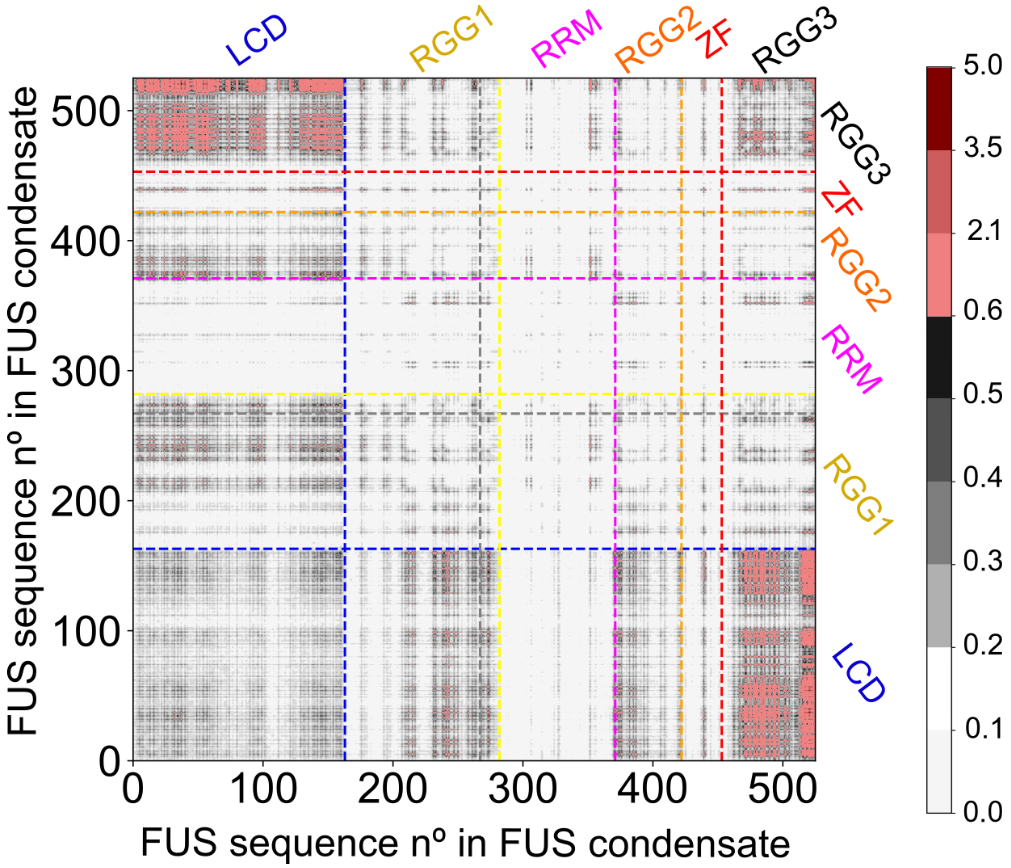


**FIG. S4:** Intermolecular contact map (expressed as residue–residue percentage of contact frequencies) of a FUS condensate.

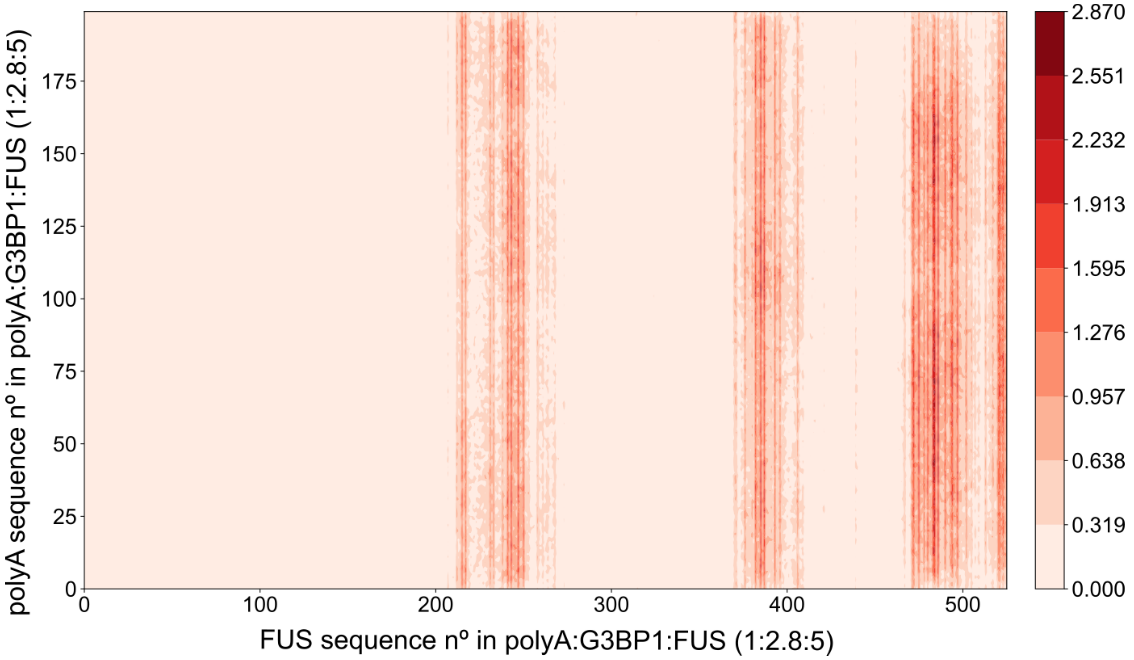


**FIG. S5:** Intermolecular contact map (expressed in residue–residue percentage of contact frequency) for polyA–FUS in polyA:G3BP1:FUS (1:2.8:5) condensate.

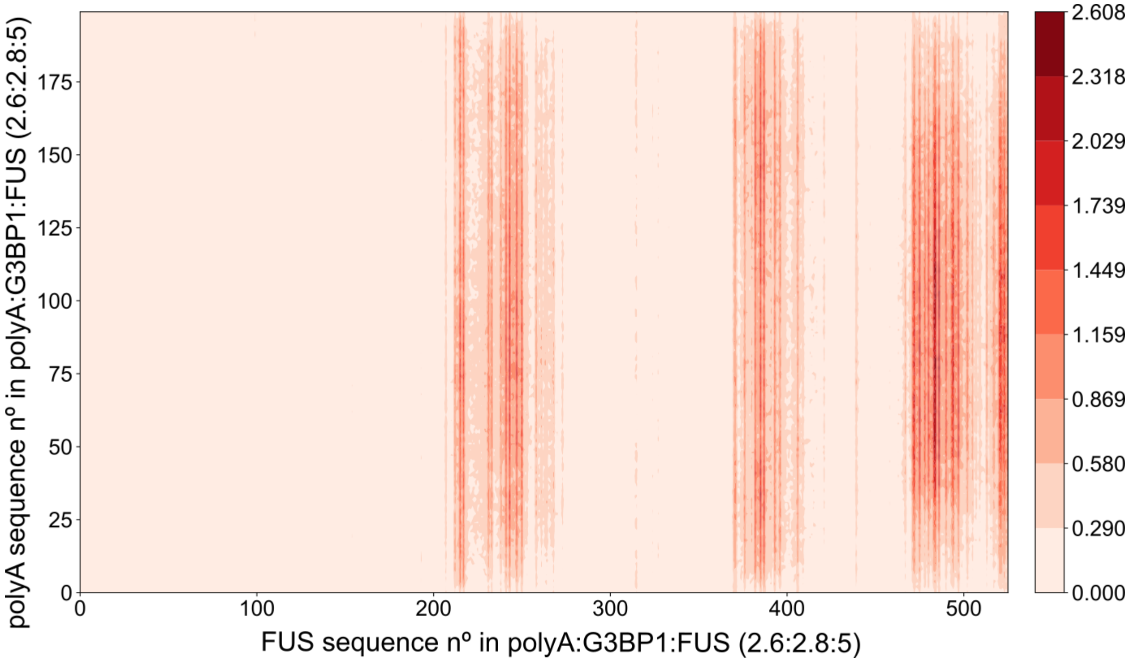


**FIG. S6:** Intermolecular contact map (expressed in residue–residue percentage of contact frequency) for polyA–FUS in polyA:G3BP1:FUS (2.6:2.8:5) condensate.

[11] J. S. Rowlinson and B. Widom, *Molecular Theory of Capillarity.* Courier Corporation, 2013.

[21] M. P. Hughes, M. R. Sawaya, D. R. Boyer, L. Goldschmidt, J. A. Rodriguez, D. Cascio, L. Chong, T. Gonen, and D. S. Eisenberg, “Atomic structures of low-complexity protein segments reveal kinked $\beta$ sheets that assemble networks,” *Science*, vol. 359, no. 6376, pp. 698–

701, 2018.
